# Up-to-date MALDI-TOF MS based identification of the complete *Corynebacterium diphtheriae* species complex for improved diagnostics

**DOI:** 10.1101/2025.07.22.665948

**Authors:** Jörg Rau, Anja Berger, Alexandra Dangel, Martin Dyk, Tobias Eisenberg, Ekkehard Hiller, Christiane Hoffmann, Peter Kutzer, An Martel, Andreas Sing, Reinhard Sting

## Abstract

The *Corynebacterium* (*C.*) *diphtheriae* species complex (*Cd*SC) comprises the closely related pathogenic species *C. diphtheriae*, *C. belfantii, C. rouxii, C. ulcerans, C. silvaticum, C. ramonii* and *C. pseudotuberculosis* according to the taxonomic revisions after 2018. Due to their close relationship, reliable species identification within the *Cd*SC is also challenging using MALDI-TOF mass spectrometry (MS) for fast and reliable identification of bacteria. However, the most commonly used commercial databases for MALDI-TOF MS do not reflect the current *Cd*SC taxonomy. Therefore, the objective of the present study is to expand the Bruker MALDI-Biotyper database version K (MBT_K_) in order to achieve reliable identification for all currently known *Corynebacterium* species within the *Cd*SC. Furthermore, the database extensions were verified with a systematic formal validation procedure in accordance with the German guideline for species identification by MALDI-TOF MS. For this, 321 strains including all seven valid species of the *Cd*SC were identified with a set of 328 spectra using the commercial MBT_K_ database and a customer-supplemented modified MBT_K_. This custom database version included 33 reference spectra of the *Cd*SC and representatives of all four recently described species. The results of the formal validations for both, the commercial and the extended database, combined partly with a single signal analysis, provide a substantiated basis for the identification of bacteria of the *Cd*SC. A key element for a fast up-to-date identification has been the exchange of user-generated sets of MSPs on the open access MALDI-UP catalogue that includes reference spectra for all members of the *Cd*SC.

**Highlights:**

- Using MALDI-TOF MS, all current species of the *C. diphtheriae* species complex are accurately identified according to the current taxonomy
- Validation was performed using spectra from different institutions and an approved gold standard protocol
- The extension and exchange of user created reference spectra via the MALDI User Platform proved to be a key tool for precise identification of bacteria

**Importance Statement:** Bacteria belonging to the *Corynebacterium* (*C.*) *diphtheriae* complex are important pathogens for humans and animals. Identification of the species of this complex described since 2018, i.e. *C. belfantii*, *C. rouxii*, *C. ramonii*, and *C. silvaticum*, using the widely approved MALDI-TOF mass spectrometry is possible. This has been achieved by improved identification using supplemented MALDI reference databases.

Comprehensive validation meets the requirements for accurate pathogen identification in accordance with recognised guidelines. This approach of identification is certainly a model for other bacterial species of interest to react on current taxonomic revisions in a short time frame.

## Introduction

Matrix-assisted laser desorption ionisation time of flight mass spectrometry (MALDI-TOF MS) is a widely used technique for the identification of microbial species in routine medical, veterinary and food microbiology (Gant & Chamot-Rooke, 2024; Thompson, 2022; Malorny et al., 2020). This method is based on the comparison of mass spectra obtained from isolates in question with the reference spectra stored in a database. Commercial databases are well organised with regard to the field of medical microbiology, but to a lesser extent in the area of food and veterinary microbiology. This applies in particular to the zoonotic pathogens of the *CdSC*, which includes the corynebacterial groups *C. diphtheriae* sensu lato (s.l.) and *C. ulcerans* s.l. and the species *C. pseudotuberculosis* which have recently undergone a revision of the species taxonomy.

*C. diphtheriae* (Lehmann & Neumann, 1896), the eponym of the species of the group *C. diphtheriae* s.l., is the causative agent of classical diphtheria of the upper respiratory tract in humans, a potentially life-threatening, highly transmissible disease (Bartlett et al., 2022; Prygiel et al. 2022). However, due to vaccination, classic diphtheria only occurs regionally in Europe (Badenschier et al., 2022; ECDC, 2022; ECDC, 2024; Hennart et al., 2023; Hoefer et al, 2025; Jaquinet et al., 2023; Walter et al., 2025). Taxonomically, *C. diphtheriae* has been subdivided into the two subspecies *diphtheriae* and *lausannense* (Tagini et al., 2018) and the biovar Belfanti has been elevated to species rank *C. belfantii* (Dazas et al., 2018). A further closely related novel *Corynebacterium* species is *C. rouxii* (Badell et al., 2020), which was formerly assigned to *C. belfantii*. This species has so far been isolated in the context of skin infections in dogs, cats and humans (Bartlett et al., 2022; Schlez et al., 2021; Sing et al., 2025).

*C. ulceran*s (Riegel et al., 1995) has already been identified as a pathogen in humans and in numerous different mammalian taxa including exotic, game, pet and livestock animals (Berger et al., 2019; Hillan et al., 2023; Eisenberg et al., 2015; Grönthal et al., 2024; Sting et al., 2023; Martel et al., 2021; Thomas et al., 2022, Wang et al., 2024). In addition to wound infections, *C. ulcerans* also causes diphtheria-like diseases of the upper respiratory tract in humans and has surpassed the incidence of *C. diphtheriae* infections in European industrialized nations since several years (ECDC, 2022). *C. ulcerans* infections in humans are mainly acquired through contact with animals, especially cats and dogs, and thus represents the most important zoonotic pathogenic agent of the *CdSC* (Hillan et al., 2023; Meinel et al., 2014; Otsuji et al., 2017; Slinko et al., 2023). *C. ulcerans* isolates from wild boars were recently assigned to the new species *C. silvaticum* (Dangel et al., 2020). This novel species has been identified as the causative agent of caseous abscesses in wild boars, Iberian pigs and, and in one reported case in a roe deer (Contzen et al., 2011; Rau et al., 2012; Eisenberg et al., 2014; Viana et al., 2023). Recently, a few cases of human infections via contact with wild boars have been reported (Berger et al., 2025). *C. ramonii* is a further novel zoonotic species of the group of *C. ulcerans* s.l. which affects humans, dogs and cats (Crestani et al., 2023; Shitada et al., 2024).

*C. pseudotuberculosis* (Eberson, 1918) is the causative agent of caseous lymphadenitis (CLA), a chronic infection mainly in sheep, goats, and camelids characterised by the formation of abscesses in lymph nodes and internal organs (Domeni, et al., 2017; Domenis et al., 2018; Sting et al., 2017; Sting et al., 2022; Hiller et al., 2024). The pathogen also causes ulcerative lymphangitis, the so-called pigeon fever in horses (Miers et al., 1980). Isolates originating from water buffaloes are the only isolates known to produce diphtheria toxin and are associated with cases of oedematous skin disease (OSD) (Selim 2001). In general, pseudotuberculosis due to *C. pseudotuberculosis* is a rare zoonosis, which is transmitted via contact with the pathogen, particularly through affected animals (Bregenzer et al., 1997; Peel et al., 1997).

The species of the *CdSC* produce the two main virulence factors, diphtheria toxin (DT; Prygiel et al., 2022; Prates et al., 2024) and phospholipase D (PLD) that are suspected to be associated with infections and exacerbation of the course of disease (Schlez et al., 2021; Aquino de Sa et al., 2013). DT is responsible for the clinical manifestations of diphtheria in humans and therefore a hallmark of pathogenicity of bacterial isolates. The presence of the underlying *tox* gene enables not only *C. diphtheriae* to produce the corresponding DT, but also some isolates of other species of the *CdSC* (Hacker et al., 2016). However, not all isolates that carry the *tox* gene express DT. PLD, a sphingomyelinase (spreading factor), has yet only been detected in *C. pseudotuberculosis*, *C. ulcerans* and *C. silvaticum* (Schlez et al. 2021, Aquino de Sa et al., 2013).

The correct identification of these closely related pathogenic corynebacteria according to the current taxonomy poses a challenge to diagnostics, especially the maintenance of an up-to-date and comprehensive MALDI-TOF MS database. Due to these obstacles commercial databases usually exhibit a temporal discrepancy in this regard. In the case of species that are relevant to human and animal health, their inclusion is of paramount importance. Rapid implementation of user generated solutions are able to close this diagnostic gap. In this context, it is important to emphasise the utmost relevance of quality assurance. The validation concepts that have been developed in accordance with national guidelines have already demonstrated their efficacy in regard to MALDI-TOF MS and Fourier-transform infrared spectroscopy (BVL, 2022; Rau et al., 2025; Oberreuter et al., 2023).

This study proposes the formal validation procedure for two databases, the commercial Bruker MALDI Biotyper database version K (MBT_K_), and an adjusted MBT_K_, which was extended with reference spectral entries from the MALDI-UP catalogue (MBT_K*_-MUP). The aim is to achieve reliable identifications of the closely related and recently described taxa of species of the *Cd*SC. This procedure is able to serve as a model for the procedure for other closely related bacterial species.

## Material and Methods

### Isolates

The type strains *C. belfantii* DSM 105776^T^, *C. diphtheriae* DSM 44123^T^, *C. pseudotuberculosis* DSM 20689^T^, *C. ramonii* CIP 112226^T^, *C. rouxii* DSM 110354^T^, and *C. ulcerans* DSM 46325^T^ were obtained from public strain collections. The type strain of *C. silvaticum* (CVUAS 4292^T^ = DSM 109167^T^) was available from the species description and previous own studies (Dangel et al., 2020; Contzen et al., 2011). In addition, the following isolates from publicly available strain collections were included: CIP 102462, CCM 1750, CCM 2823 (assigned as *C. ulcerans*), DSM 43988, ATCC 13812 = DSM 43989 (*C. diphtheriae*), and DSM 7177 (*C. pseudotuberculosis)*. In addition to the *Cd*SC strains from public strain collections, further 308 *Cd*SC isolates were used for the generation of reference spectra and/or for validation purposes (Table S1).

Except for one isolate from a horse, all utilised isolates of *C. diphtheriae* s.s. (n=26), and *C. belfantii* (n=6) originated from humans with clinical cases that had been sent to the German National Consiliar Laboratory for Diphtheria (LGL Bayern). Isolates of *C. ulcerans* s.s. were obtained from humans (n=18), and from ten different animal species (n=83). The previously known six own *C. rouxii* isolates were obtained from canids and *C. silvaticum* isolates (n = 86) were cultured from diseased wild boars and one roe deer (Schlez et al., 2021; Rau et al., 2019). Two recent *C. silvaticum* isolates were of human origin (Berger et al., 2025). Except for the type strain of *C. ramonii,* no further assigned isolate of this species was available at the time of the start of this study (Table S1).

For testing the exclusivity of the method 1,264 isolates outside the *Cd*SC were used, representing 844 species of 218 genera, including 127 bacterial isolates from other members of the phylum Actinomycetota (Actinobacteria) (cg. Nouioui et al., 2018). In this way, a broad taxonomic selection of microorganisms were included for the validation procedure (Table S1).

The human isolates included in this study had been confirmed by the National Consiliar Laboratory for Diphtheria (LGL Bayern) by MALDI-TOF MS based species identification (Bruker microflex with MBT_K_ database) and a *tox* gene PCR, followed by a modified ELEK test demonstrating DT expression if the PCR result was positive (Dangel et al., 2020; Engler et al., 1997).

The production of PLD was detected by means of two tests for the CAMP phenomenon (inhibition of the outer haemolysis zone of an orthogonal growing *Staphylococcus aureus* isolate and synergistic enhancement of the haemolysis with orthogonal growing *Rhodococcus hoagii*) (cf. Rau et al., 2019; Barksdale et al., 1981).

The confirmation of potentially *C. ramonii* isolates was achieved by whole genome DNA sequencing (WGS), followed by an average nucleotide identity analysis using PyANI, an open-source python-based tool for calculating Average Nucleotide Identity (ANI) software (v. 0.2.12, https://pyani.readthedocs.io). WGS was performed on a NovaSeq 6000 (S4 Reagent Kit v1.5) with 2x150 bp paired-end reads subsequent to Nextera XT library preparation (both Illumina, San Diego, USA), using 1 ng of genomic DNA. Library preparation and sequencing were carried out by CeGaT GmbH (Tübingen, Germany). The sequencing data sets were assembled with the AQUAMIS pipeline (Deneke et al., 2021). The genomes were compared to a selection of publicly available reference genomes in the NCBI (https://www.ncbi.nlm.nih.gov/), downloaded on 04.09.2024. The complete list of sequence data sets used can be found in the appendix (Table S2). These data sets included, for example the *C. pseudotuberculosis* reference genome of strain MEX29 (NZ_CP016826.1) as well as *C. belfantii* FRC0043 (GCF_900205605), *C. diphtheriae* ISS3319 (NZ_CP0252091), *C. rouxii* FRC0190 (NZ_LR738855), *C. ulcerans* 809 (NC_017317), *C. ramonii* FRC0011 (GCF_000767685.1) and *C. silvaticum* PO100/5 (NZ_CP021417). ANIm (MUMmer algorithm) and ANIb (BLASTN+ algorithm) analyses were performed with help of PyANI as previously described by Hiller et al. (2024) (Figure S6). An ANI match with more than 95% identity to a reference isolate was used to determine the species (Goris et al., 2007).

All individual available results of characterizing tests for all isolates used are depicted in Table S1.

### Cultivation and sample preparation

Corynebacteria and other microorganisms belonging to different taxonomic groups were cultivated in accordance with their specific requirements. The individual growth conditions of all bacteria utilised for single spectra acquisitions are accessible through the MALDI-UP (MUP) catalogue (Rau et al., 2016). A selection of these data is presented in Table S1. The ethanol-formiate extraction protocol was used for generating MALDI-TOF mass reference spectra as recommended by the manufacturer. In addition, the direct transfer and extended direct transfer protocols were applied to record the set of individual spectra used for validation. These latter protocols correspond to those used in the routine workflow in many laboratories (Table S1) (Pranada et al., 2016; Kostrzewa & Maier, 2017; Rau et al., 2025). In addition, spectra listed in MUP were included, which were created using a special sample preparation procedure for highly pathogenic bacteria (Lasch et al., 2008).

### MALDI-TOF MS – Databases

Two databases were used and the results were compared.

#### MBT_K_

The commercial database version K for the Bruker MALDI-Biotyper Compass system (MBT_K_) was released in 2022 as a “research use only” (RUO) application (Bruker Daltonik, Bremen, Germany). This database includes 11,897 reference entries, deposited as so called main spectra or main spectra projections (MSP) by the manufacturer, which were derived from 4,320 bacterial species (database product information, Bruker, 2022). In the MBT_K_ database, only three species of the *Cd*SC have been included up to the year 2018: *C. diphtheriae*, *C. ulcerans* and *C. pseudotuberculosis*. For these entries no subspecies information was given by the manufacturer (Table 1). Due to lacking specific MALDI-TOF reference mass spectra, the current Bruker MBT_K_ database is not able to differentiate the species of *C. diphtheriae* s.l. and *C. ulcerans* s.l. that have been described since 2018.

**Table 1.** MALDI-TOF MS reference spectra of *Corynebacterium diphtheriae* species complex bacteria (*Cd*SC), available in the MALDI Biotyper database version K (MBT_K_), and additional reference spectra listed on MALDI User Platform (MUP) used in this study. MBT_K_*: Commercial database modified by extraction of entries of unclear assignment with regard to the new species of the *Cd*SC. MUP: Custom made database entries selected from the MALDI-UP catalogue. The MUP catalogue number is given for user made entries. T: type strain; bv: biovar; gv: genomovar.

| Species | Isolate/<br>strain |  | Reference entry name (MSP) | Database<br>source | Database |  |
| --- | --- | --- | --- | --- | --- | --- |
|  |  |  |  |  | MBT<sub>K</sub> | MBT<sub>K</sub>*<br>+MUP |
| *C. belfantii* | DSM 105776 | T | *Corynebacterium belfantii* DSM 105776 CVUAS | MUP 1781 |  | X |
|  | KL0171 |  | *Corynebacterium belfantii* KL0171 CVUAS | MUP 1745 |  | X |
|  | KL0236 |  | *Corynebacterium belfantii* KL0236 CVUAS | MUP 1746 |  | X |
| *C. diphtheriae* | DSM 44123 | T | *Corynebacterium diphtheriae* DSM 44123<sup>T</sup> DSM_2 | MBT<sub>K</sub> | X |  |
|  |  |  | *Corynebacterium diphtheriae diphtheriae* DSM 44123 CVUAS | MUP 6620 |  | X |
|  | CCUG 15935 |  | *Corynebacterium diphtheriae* CCUG 15935 CCUG | MBT<sub>K</sub> | X |  |
|  | CCUG 55537 |  | *Corynebacterium diphtheriae* CCUG 55537 CCUG | MBT<sub>K</sub> | X |  |
|  | DSM 43988 |  | *Corynebacterium diphtheriae* DSM 43988 DSM | MBT<sub>K</sub> | X |  |
|  | 1G12072319_3e |  | *Corynebacterium diphtheriae* 1G12072319_3e MVD | MBT<sub>K</sub> | X |  |
|  | 44_R9 |  | *Corynebacterium diphtheriae* 44_R9 ISB | MBT<sub>K</sub> | X |  |
|  | 50_R5 |  | *Corynebacterium diphtheriae* 50_R5 ISB | MBT<sub>K</sub> | X |  |
|  | R1_36_29 |  | *Corynebacterium diphtheriae* R1_36_29 ISB | MBT<sub>K</sub> | X |  |
|  | KL0163 |  | *Corynebacterium diphtheriae gravis* KL0163 CVUAS | MUP 1747 |  | X |
|  | KL0167 |  | *Corynebacterium diphtheriae gravis* KL0167 CVUAS | MUP 1748 |  | X |
|  | KL0187 |  | *Corynebacterium diphtheriae gravis* KL0187 CVUAS | MUP 1749 |  | X |
|  | KL0232 |  | *Corynebacterium diphtheriae gravis* KL0232 CVUAS | MUP 1750 |  |  |
|  | KL0237 |  | *Corynebacterium diphtheriae gravis* KL0237 CVUAS | MUP 1751 |  | X |
|  | KL0240 |  | *Corynebacterium diphtheriae gravis* KL0240 CVUAS | MUP 1752 |  | X |
|  | KL0173 |  | *Corynebacterium diphtheriae mitis* KL0173 CVUAS | MUP 1753 |  | X |
|  | KL0179 |  | *Corynebacterium diphtheriae mitis* KL0179 CVUAS | MUP 1754 |  | X |
|  | KL0235 |  | *Corynebacterium diphtheriae mitis* KL0235 CVUAS | MUP 1755 |  | X |
| *C. pseudotuberculosis* | DSM 20689 | T | *Corynebacterium pseudotuberculosis* DSM 20689<sup>T</sup> DSM | MBT<sub>K</sub> | X | X |
|  |  |  | *Corynebacterium pseudotuberculosis* Ovis gv ovis DSM 20689 CVUAS | MUP 7393 |  | x |
|  | 59_D6_coll |  | *Corynebacterium pseudotuberculosis* 59_D6_coll ISB | MBT<sub>K</sub> | x | x |
|  | 62_D6_coll |  | *Corynebacterium pseudotuberculosis* 62_D6_coll ISB | MBT<sub>K</sub> | X | X |
|  | 64_D6_coll |  | *Corynebacterium pseudotuberculosis* 64_D6_coll ISB | MBT<sub>K</sub> | X | X |
|  | GD7 |  | *Corynebacterium pseudotuberculosis* GD7 GDD | MBT<sub>K</sub> | X | X |
|  | GD8 |  | *Corynebacterium pseudotuberculosis* GD8 GDD | MBT<sub>K</sub> | X | X |
|  | X x 102968 C |  | *Corynebacterium pseudotuberculosis* x_x_102968_C IBS | MBT<sub>K</sub> | X | X |
|  | 992 |  | *Corynebacterium pseudotuberculosis* Equi NTTB 992 CVUAS | MUP 935 |  | X |
|  | CVUAS 32689 |  | *Corynebacterium pseudotuberculosis* Ovis gv-camelid CVUAS 32689 CVUAS | MUP 2398 |  | X |
|  | CVUAS 5583,2 |  | *Corynebacterium pseudotuberculosis* bv Ovis gv-camelid CVUAS 5583.2 CVUAS | MUP 958 |  | X |
| *C. ramonii* | CIP 112226 | T | *Corynebacterium ramonii* CIP 112226<sup>T</sup> CVUAS | MUP 6614 |  | X |
|  | CVUAS 34267 |  | *Corynebacterium ramonii* CVUAS 34267 CVUAS | MUP 7345 |  | X |
|  | C104 |  | *Corynebacterium ramonii* C104 CVUAS | MUP 7346 |  | X |
| *C. rouxii* | DSM 110354 | T | *Corynebacterium rouxii* DSM 110354 CVUAS | MUP 1782 |  | X |
|  | 191012535 |  | *Corynebacterium rouxii* 191012535 CVUAS | MUP 1935 |  | X |
|  |  |  | *Corynebacterium rouxii* 191012535_LHL-GI | MUP 1744 |  | X |
|  | 211002017-1 |  | *Corynebacterium rouxii* 211002017-1_LHL-GI | MUP 3235 |  | X |
|  | 211002017-2 |  | *Corynebacterium rouxii* 211002017-2_LHL-GI | MUP 3236 |  | X |
|  | C138 |  | *Corynebacterium rouxii* C138 CVUAS | MUP 813 |  |  |
|  | CVUAS 3559,2 |  | *Corynebacterium rouxii* CVUAS 3559,2 CVUAS | MUP 1014 |  | X |
|  | KL1306 |  | *Corynebacterium rouxii* KL 1306 CVUAS | MUP 1783 |  |  |
|  | KL1663 |  | *Corynebacterium rouxii* KL 1663 CVUAS | MUP 1901 |  |  |
| *C. silvaticum* | CVUAS 4292 = DSM 109166 | T | *Corynebacterium silvaticum* CVUAS 4292 CVUAS | MUP 0083 |  | X |
|  | CVUAS 6455 |  | *Corynebacterium silvaticum* CVUAS 6455 CVUAS | MUP 0922 |  | X |
| *C. ulcerans* | DSM 46325 | T | *Corynebacterium ulcerans* DSM 46325<sup>T</sup> DSM | MBT<sub>K</sub> | X |  |
|  |  |  | *Corynebacterium ulcerans* DSM 46325 CVUAS | MUP 2212 |  | X |
|  | 66_D6_coll |  | *Corynebacterium ulcerans* 66_D6_coll ISB | MBT<sub>K</sub> | X |  |
|  | 67_D6_coll |  | *Corynebacterium ulcerans* 67_D6_coll ISB | MBT<sub>K</sub> | X |  |
|  | 71_D6_coll |  | *Corynebacterium ulcerans* 71_D6_coll ISB | MBT<sub>K</sub> | X |  |
|  | MCW 10006 |  | *Corynebacterium ulcerans* MCW_10006 MCW | MBT<sub>K</sub> | X |  |
|  | VA22435_07 |  | *Corynebacterium ulcerans* VA22435_07 ERL | MBT<sub>K</sub> | X |  |
|  | 131011719 |  | *Corynebacterium ulcerans* NT 131011719 CVUAS | MUP 1015 |  | X |
|  | 141001548 |  | *Corynebacterium ulcerans* NT 141001548 CVUAS | MUP 1021 |  | X |
|  | 07UVF148 |  | *Corynebacterium ulcerans* 07UVF148 LLBB | MUP 0966 |  | X |
|  | CVUAS 10306 |  | *Corynebacterium ulcerans* TTB CVUAS 10306 CVUAS | MUP 0945 |  | X |
|  | S-477-6-21 |  | *Corynebacterium ulcerans* S-477-6-21 CVUAS | MUP 2339 |  | X |

#### MBT_K_*+MUP

In order to be able to identify the new species of the *Cd*SC, a revision of the database MBT_K_ was required. For this purpose, 33 custom made MSPs from the MUP catalogue for all current seven *Cd*SC species were created by three laboratories in order to replace the previous MSPs of *C. diphtheriae* (s.l.) and *C. ulcerans* (s.l.) in the commercial reference database. These user made entries were integrated into the greater MUP-database using the project section of the Biotyper Compass Explorer software module (Bruker) (Table 1). Four further MSPs were used in the dendrogram analysis (Figure 1). Reference spectra measurements and calculations of MSPs followed the manufacturer’s manual and commentary (Kostrzewa & Maier, 2017).

**Figure 1.**
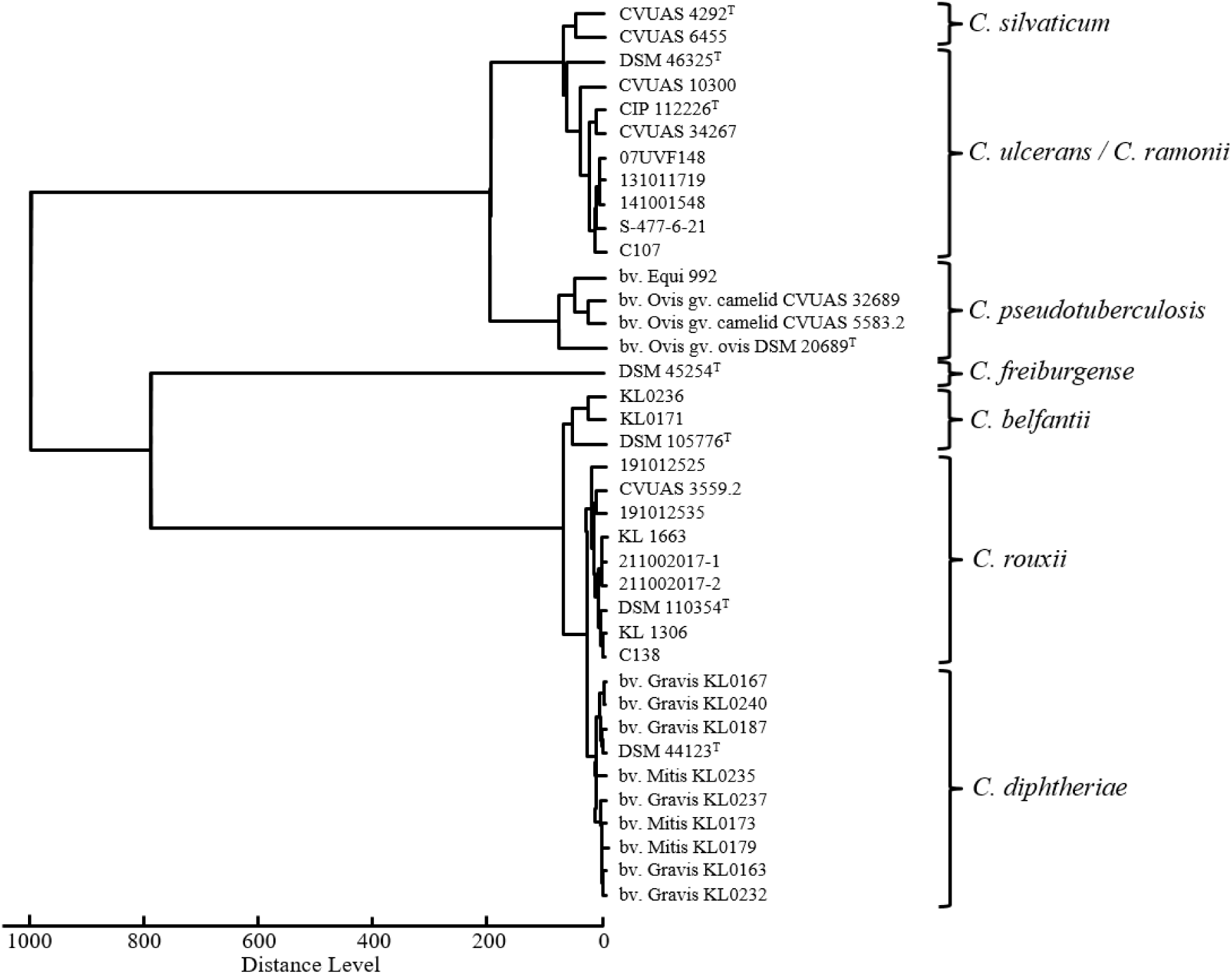
Dendrogram of selected MALDI-TOF reference mass-spectra from all current species of the *Corynebacterium diphtheriae* species complex, created by MALDI-UP users, is visualising the spectral relationships of the species.

These procedures were recently described in more detail (Rau et al., 2025).

#### Database cross check

As the taxonomic species delineated since 2018 have not yet been included in the commercial version of the MBT_K_, it can be assumed that individual reference isolates in the current database are still not recognized to belong to one of the newly described species. To confirm this suspicion, a “MSP cross-check” was conducted. Every single MSP of the commercial MBT_K_ database was identified using the MUP database. The results were evaluated using the first two hits obtained, analogous to the evaluation of individual spectra. This procedure shows MSPs of the commercial database, whose taxonomic status may be outdated.

### Validation spectra

For validation purposes, a total of 1,609 single MALDI-TOF mass spectra obtained from 1,585 isolates were selected from the spectra listed in the MUP catalogue (version 05.03.2025, https://maldi-up.ua-bw.de) (cf. Rau et al., 2016). Of these, 457 spectra from 448 isolates belonged to the phylum Actinomycetota, including 328 spectra (321 isolates) from *Cd*SC members. The spectra used originated from pathogenic bacterial species belonging to the phylum Actinomycetota, e. g. *Actinomyces* (n=5 species), *Arcanobacterium* (n=11),

*Schaalia* (n=5) or other *Corynebacterium* spp. (n=37). Further information on every individual spectrum and the custom reference spectra is provided in MUP. This catalogue includes metadata of the isolates, cultivation conditions, the method of sample preparation used for creating MALDI-TOF mass spectra, the used MALDI-TOF MS system, and the user creating the spectra (see Rau et al, 2016; Table S1). Hence, spectra from 17 laboratories were used for the complete set of spectra obtained from MUP.

### Validation procedure

The validation procedure has recently been described (Rau et al., 2025) and follows the guidelines created by the working group MALDI-TOF, pursuant to §64 of the German Food and Feed Code (BVL, 2022; Rau et al., 2022). These guidelines, which are used for MALDI-TOF MS species identification in the food sector, have been adopted unchanged for the differentiation of *Cd*SC bacteria. In order to examine the effect of custom-made database additions on the identification results, two databases were compared using the same validation spectra set. The first was the latest commercial MBT_K_ database, and the second was a modified MBT_K_, with an exchange of the commercial *CdS*C reference entries from *C. diphtheriae* (s.l.) and *C. ulcerans* (s.l.) with 33 custom-made reference MSPs of the *Cd*SC obtained from MALDI-UP (Table 1). This version of the custom database includes 1,486 user-made entries of a broad number of bacterial species (Table 1).

Identification results were scored according to the manufacturer’s instructions (Bruker). The first result in the list has to yield a score value of at least 2.0, and the second result has to show no contradictory species, that exceeds a score value of ≥2.0. This straightforward rule was applied to the same extensive spectra set (n=1,609) for both database combinations. The detailed results are presented as reports in the supplementary Table S3a-S3i for the commercial MBT_K_ database and Table S4a-S4f for the modified and extended database MBT_K_*+MUP.

### Manual analysis of spectra

Crestani and colleagues observed a specific MALDI-TOF mass spectrum signal at 5,405.4 *m/z* distinguishing the newly described species *C. ramonii* from *C. ulcerans* s.s. (Crestani et al., 2023). Supplementary Figures 2a and 2b present overlays of mass spectra from all *C. ulcerans* s.s. and *C. ramonii* isolates of this study, confirming the *m/z* (5,405.4 +/- 2.0) window in which the specific signal of *C. ramonii* is recognised. Therefore, raw spectra were loaded into the FlexAnalysis program (version 3.4, Bruker) for manual analysis. Baseline correction and smoothing were performed using settings consistent with the MSP preparation procedure (MBT standard method, Bruker). Peak maxima of interest were manually assigned (Figures 2a and 2b).

**Figure 2a.**
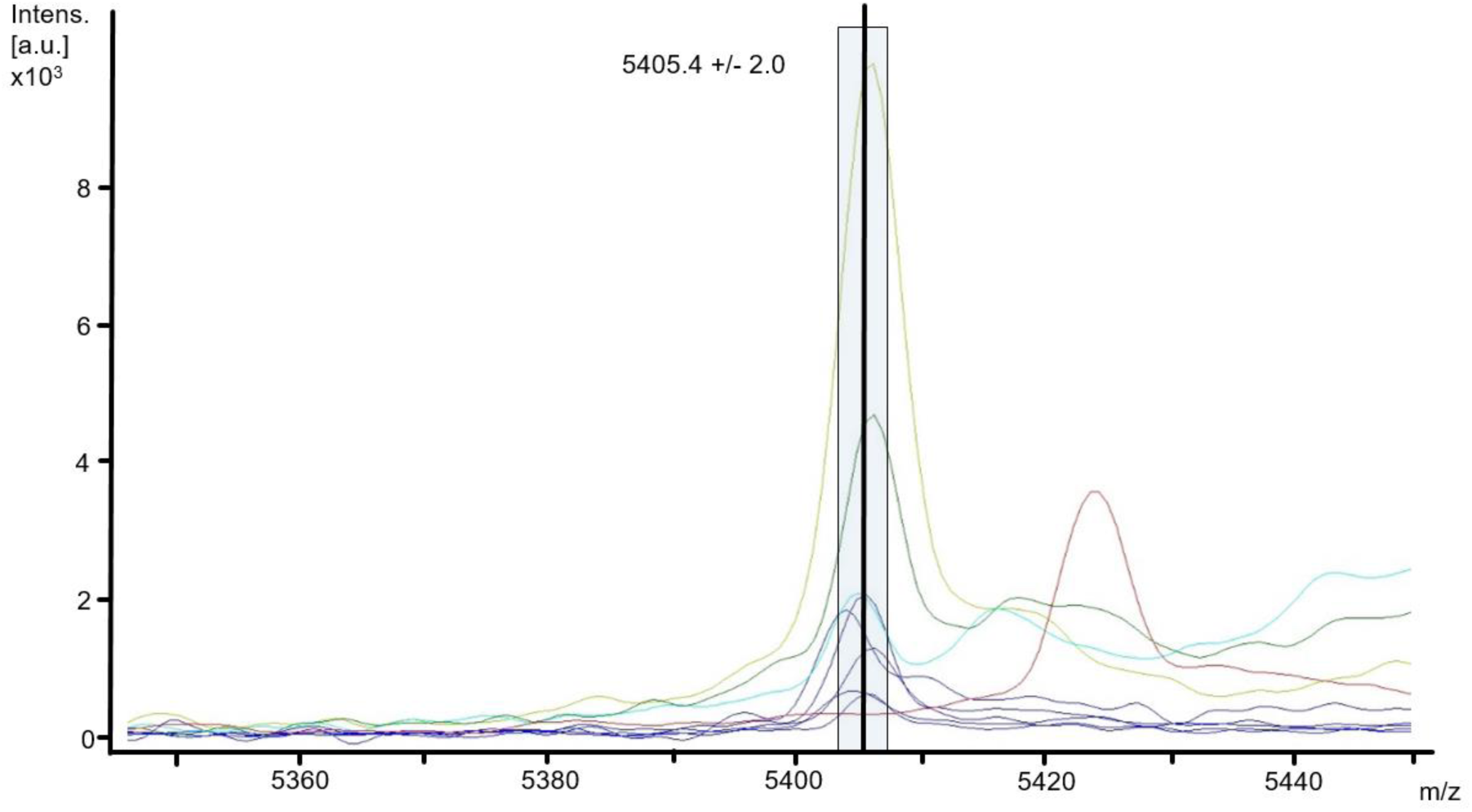
Section of the MALDI-TOF mass spectra, ranging from 5,350 – 5,450 *m/z*, displaying the signal characteristic of *Corynebacterium* (*C.*) *ramonii* at 5405.4 *m/z*, as outlined by Crestani et al., 2023. green: *C. ramonii* CIP 112226^T^, red: *C. ulcerans* DSM 46325^T^, dark blue: spectra of *C. ramonii* isolates C86, C104, C139, C167, CVUAS 34267, all from hedgehogs, and pale blue and light green: KL3299A and CCM 2825 from humans (for metadata of the isolates, see Supplementary Table S1). The bar shows the range of +/-2 *m/z* around the specific signal. ^T^: type strain.

**Figure 2b.**
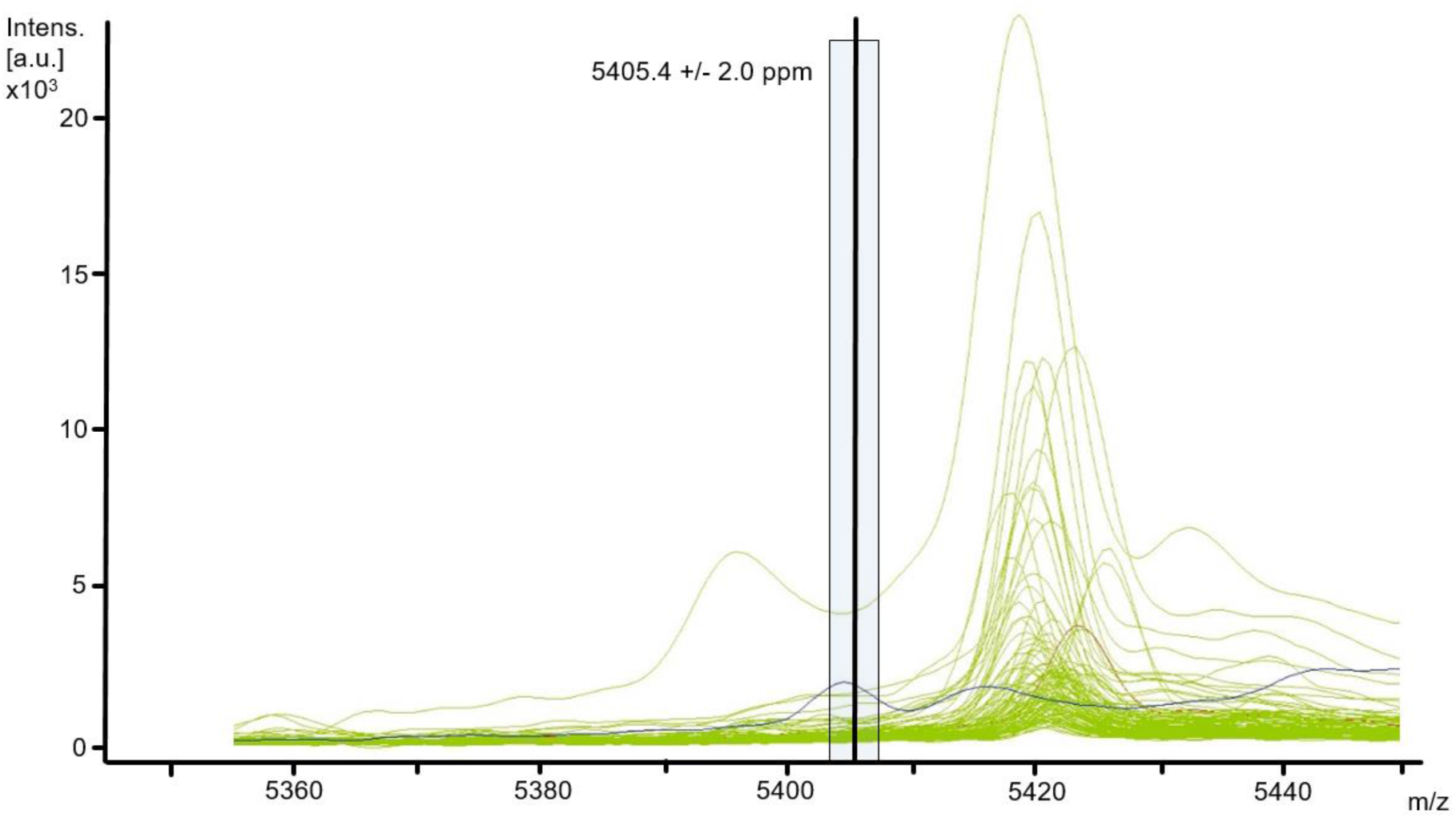
Spectral *m/z* window according to Figure 2a. blue: *Corynebacterium* (*C.*) *ramonii* CIP 112226^T^, green: overlay of single spectra of 103 *C. ulcerans* s.s. isolates, including in red the type strain of *C. ulcerans* DSM 46325^T^ (for metadata of the isolates, see Supplementary Table 1). The bar shows the range of +/-2 *m/z* around the signal specific for *C. ramonii* according Crestani et al., 2023. ^T^: type strain.

## Results and Discussion

Numerous bacteria are currently facing significant revisions in taxonomy, promoted by a strong increase in data from WGS. Therefore, it is essential to align the key tools for species identification like MALDI-TOF MS to current revisions in order to ensure reliable and up–to-date species level identifications. This is particularly the case for microorganisms of medical and veterinary relevance, such as members of the *Cd*SC (Badell et al., 2020; Dangel et al., 2020; Dazas et al., 2018; Crestani et al., 2023). *Cd*SC pathogens cause serious diseases in humans and animals, the most prominent being diphtheria, which is therefore subject to a strict disease notification obligation in the EU and therefore requires reliable diagnosis (ECDC, 2024; EU-Commission, 2018; Hoefer et al., 2025). Detailed information on the known combinations of *Cd*SC species, clinical pictures and affected host species is summarised in Table S5.

MALDI-TOF MS is a tool that has achieved widespread acceptance and has become the method of choice for routine identification of bacteria in both medical and veterinary diagnostics (Malorny et al., 2020; Timofte et al., 2023; Gant & Chamot-Rooke, 2024; Thompson, 2022). This technique makes it possible to keep up regularly with new taxonomic developments. This requires prompt database updates by implementation of supplementary or revised reference spectra (see Rau et al., 2025). In a first step, a comprehensive database is necessary that includes a sufficient number of reference spectra obtained from the target organisms. Subsequently, a formal validation procedure for the new parameters in focus, as well as for non-target parameters is necessary for a valid and sustainable expansion and update of the used database.

### Results with the commercial MBT_K_ database

The commercial MALDI Biotyper database MBT_K_ contains reference spectra only for three representatives of the *Cd*SC (n=21), including entries for the type strains of *C. pseudotuberculosis*, *C. ulcerans*, and *C. diphtheriae* (Table 1), reflecting the taxonomic status as reviewed by Oliveira et al., 2017. However, the species described since 2018 have not yet been included in this reference dataset (Bruker MBT_K_). With the aim of including the new species of the *Cd*SC for improving the species identification accuracy we created and exchanged 33 custom made reference entries to extent the commercial MBT_K_ database (Table 1; Figure 1).

Thus, using the commercial MBT_K_ among the validation spectra set, spectra of the current taxonomic species *C. diphtheriae* s.s. (n=31), *C. belfantii* (n=7) and *C. rouxii* (n=7) yielded consistently score values greater than 2.0 and were consequently supposedly assigned to “*C. diphtheriae”*. From this spectra set, one *C. rouxii* isolate could not be identified. Considering the current taxonomy, *C. belfantii* and *C. rouxii* were not correctly identified as separate species by the MBT_K_. Therefore, using the commercial MBT_K_ only allows identification at the level of *C. diphtheriae* s.l..

Shortcomings in the accurate identification of the corynebacterial species *C. ulcerans* and *C. silvaticum* also occur when using both, the former commercial MBT_E_ database (7,854 entries, Bruker, 2018) or the later MBT_K_ database (Rau et al., 2019). In addition, the recently described species *C. ramonii* cannot be identified by the current commercial MBT_K_ database (Crestani et al., 2023). In consequence, all validation spectra of *C. ulcerans* s.s. (n=105), *C. silvaticum* (n=88), and *C. ramonii* (n=9) were identified as *C. ulcerans* s.l. by the commercial MBT_K_ (Table 2). In contrast, all confirmed *C. pseudotuberculosis* isolates (n=80) were correctly identified using the commercial MBT_K_ with a mean score value of >2.0 (Table 2).

**Table 2.** Species identifications of *Corynebacterium diphtheriae* species complex bacteria samples by MALDI-TOF MS using the commercial database MBT_K_. True/False: the species was correctly/not correctly identified. All samples within the control group did not belong to the parameter of interest. Individual results for any sample were given in Supplementary Tables S2a-S2i. \**C. belfantii* and *C. rouxii* are not represented in the database used / \*\**C. ramonii* and *C. silvaticum* are not represented in the database used.

| Species / Parameter of interest | Samples | Score mean | Score STD | Samples identified | Identification rate (%) | True positive | False negative | True positive rate (%) of identified samples | False negative rate (%) of identified samples | Samples of the control group | Samples identified | Identification rate (%) | True negative | False positive | True negative rate (%) of identified samples | False positive rate (%) of identified samples | Criteria of the BVL guideline fulfilled |
| --- | --- | --- | --- | --- | --- | --- | --- | --- | --- | --- | --- | --- | --- | --- | --- | --- | --- |
| ***C. diphtheriae* s.l.** |  |  |  |  |  |  |  |  |  |  |  |  |  |  |  |  |  |
| *C. belfantii* | 7 | 2.19 | 0.09 | 7 | 100 | 0 | 7* | 0 | 100* | 1602 | 803 | 50.1 | 803 | 0 | 100 | 0 | no |
| *C. diphtheriae* s.s. | 31 | 2.41 | 0.11 | 31 | 100 | 31 | 0 | 100 | 0 | 1578 | 786 | 49.8 | 772 | 14* | 98.2 | 1.8* | no |
| *C. rouxii* | 8 | 2.12 | 0.11 | 7 | 87.5 | 0 | 7* | 0 | 100* | 1601 | 803 | 50.2 | 803 | 0 | 100 | 0 | no |
| *C. diphtheriae* s.s., *C. belfantii*, *C. rouxii* interpreted as ***C. diphtheriae* s.l.** | 46 | 2.33 | 0.16 | 45 | 97.8 | 45 | 0 | 100 | 0 | 1563 | 772 | 49.4 | 772 | 0 | 100 | 0 | yes* |
| ***C. pseudotuberculosis*** | 80 | 2.36 | 0.17 | 80 | 100 | 80 | 0 | 100 | 0 | 1529 | 723 | 47.3 | 723 | 0 | 100 | 0 | yes |
| ***C. ulcerans* s.l.** |  |  |  |  |  |  |  |  |  |  |  |  |  |  |  |  |  |
| *C. silvaticum* | 88 | 2.10 | 0.10 | 78 | 88.6 | 0 | 78** | 0 | 78** | 1521 | 803 | 52.8 | 803 | 0 | 100 | 0 | no |
| *C. ramonii* | 9 | 2.37 | 0.09 | 9 | 100 | 0 | 9 | 0 | 100** | 1600 | 803 | 50.2 | 803 | 0 | 100 | 0 | no |
| *C. ulcerans* s.s. | 105 | 2.37 | 0.10 | 105 | 100 | 105 | 0 | 200 | 0 | 1504 | 785 | 52.2 | 698 | 87* | 88.9 | 11.1** | no |
| *C. ulcerans* s.s., *C. ramonii*, *C. silvaticum* interpreted as ***C. ulcerans* s.l.** | 202 | 2.25 | 0.17 | 202 | 100 | 202 | 0 | 100 | 0 | 1485 | 698 | 46.7 | 698 | 0 | 100 | 0 | yes** |

Corynebacteria not belonging to the *Cd*SC and other species of the phylum Actinomycetota as well as more distantly related bacteria included in the validation set, comprising more than 1,500 individual spectra, were clearly separated from the *Cd*SC species using the commercial MBT_K_ database (Table 2).

In summary, the commercial MBT_K_ database allows the identification of the *Cd*SC species only for the three species *C. diphtheriae*, *C. ulcerans*, and *C. pseudotuberculosis* according the taxonomy as of 2018, but does not differentiate the currently recognised and medically relevant species *C. belfantii, C. ramonii, C. rouxii,* and *C. silvaticum*.

However, the commercial MBT_K_ provides the “matching hint” “*Species pseudotuberculosis / ulcerans of the genus Corynebacterium have very similar patterns: Therefore distinguishing this species is difficult”* for *C. pseudotuberculosis* and *C. ulcerans* (Bruker, 2022). Based on the results of the validation study presented here, the previous matching hint for the MBT_K_ may be replaced by more specific advice in each case (Table 3).

**Table 3.** Matching hints of *Corynebacterium diphtheriae s*pecies complex samples by MALDI-TOF MS for identifications results with MBT_K_. Utilising solely the commercial MBT_K_ database version, the following matching hints for the identification were proposed. Because of the lack of specific MSPs for certain species in the database used, a differentiation within the mentioned sensu lato groups is not applicable.

| Species identified by MBT<sub>K</sub> | Proposed matching hint to avoid misidentification |
| --- | --- |
| *C. diphtheriae* | This result should be interpreted as *C. diphtheriae* s.l., comprising *C. diphtheriae* s.s., *C. belfantii*, and *C. rouxii* |
| *C. pseudotuberculosis* | No matching hint |
| *C. ulcerans* | This result should be interpreted as *C. ulcerans* s.l., comprising *C. ulcerans* s.s., *C. silvaticum*, and *C. ramonii* |

### Cross check of database versions and single signal test for *C. ramonii*

Due to the uncertain species identification of the members of the *Cd*SC according to the taxonomy after 2018, MSPs for the three species were extracted from the MBT_K_ and identified using the comprehensive self-created reference spectra set. However, the identification results obtained in this way were not always consistent (Table 4). Thus, among the species attributable to *C. diphtheriae* s.l. seven of eight MSPs showed a matching result for *C. diphtheriae* s.s., consistent in the first and second hit. In contrast, the MSP “*Corynebacterium diphtheriae* 44_R9 ISB” was assigned as *C. belfantii,* the former member of *C. diphtheriae.* Similarly, analysis of the six MBT_K_-MSPs of *C. ulcerans* s.l. resulted in several hits for the type strain of *C. ramonii* with a score of ≥2.2 as the first and/or the second hit result (Table 5). In contrast, no conflicts in the assignment could be identified for the still uniform species *C. pseudotuberculosis*.

**Table 4.**
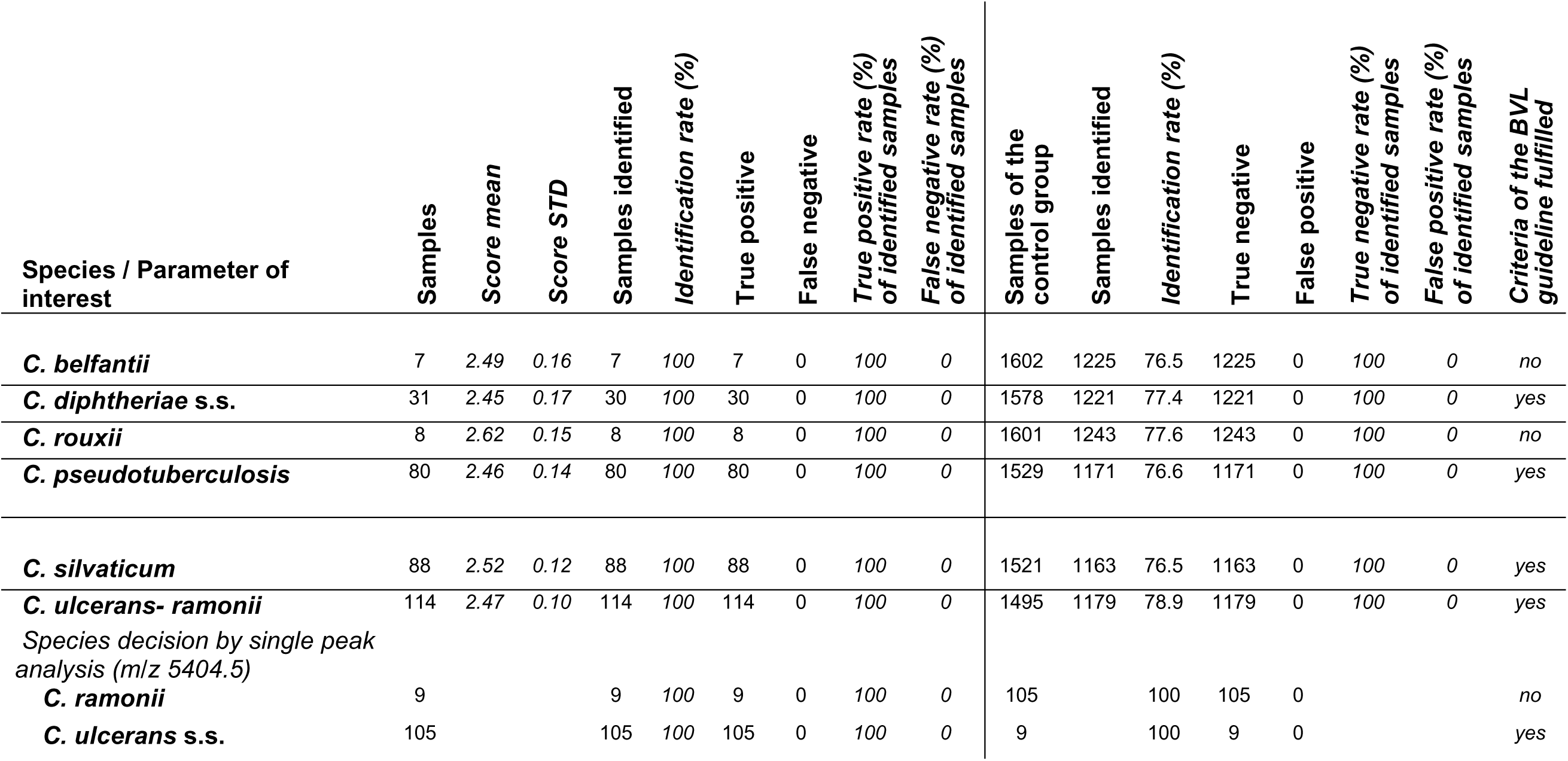
Results of species identification of *Corynebacterium diphtheriae s*pecies complex samples by MALDI-TOF MS using the modified MBT_K_*, extended with the MALDI-UP database selection (MBT_K_*+MUP). True/False: the species was correctly/not correctly identified. All samples within the control group did not belong to the parameter of interest. Individual results for any sample were given in Supplementary Tables S3a-S3i.

| Species / Parameter of interest | Samples | Score mean | Score STD | Samples identified | Identification rate (%) | True positive | False negative | True positive rate (%) of identified samples | False negative rate (%) of identified samples | Samples of the control group | Samples identified | Identification rate (%) | True negative | False positive | True negative rate (%) of identified samples | False positive rate (%) of identified samples | Criteria of the BVL guideline fulfilled |
| --- | --- | --- | --- | --- | --- | --- | --- | --- | --- | --- | --- | --- | --- | --- | --- | --- | --- |
| ***C. belfantii*** | 7 | 2.49 | 0.16 | 7 | 100 | 7 | 0 | 100 | 0 | 1602 | 1225 | 76.5 | 1225 | 0 | 100 | 0 | no |
| ***C. diphtheriae* s.s.** | 31 | 2.45 | 0.17 | 30 | 100 | 30 | 0 | 100 | 0 | 1578 | 1221 | 77.4 | 1221 | 0 | 100 | 0 | yes |
| ***C. rouxii*** | 8 | 2.62 | 0.15 | 8 | 100 | 8 | 0 | 100 | 0 | 1601 | 1243 | 77.6 | 1243 | 0 | 100 | 0 | no |
| ***C. pseudotuberculosis*** | 80 | 2.46 | 0.14 | 80 | 100 | 80 | 0 | 100 | 0 | 1529 | 1171 | 76.6 | 1171 | 0 | 100 | 0 | yes |
| ***C. silvaticum*** | 88 | 2.52 | 0.12 | 88 | 100 | 88 | 0 | 100 | 0 | 1521 | 1163 | 76.5 | 1163 | 0 | 100 | 0 | yes |
| ***C. ulcerans- ramonii*** | 114 | 2.47 | 0.10 | 114 | 100 | 114 | 0 | 100 | 0 | 1495 | 1179 | 78.9 | 1179 | 0 | 100 | 0 | yes |
| *Species decision by single peak analysis (m/z 5404.5)* |  |  |  |  |  |  |  |  |  |  |  |  |  |  |  |  |  |
| ***C. ramonii*** | 9 |  |  | 9 | 100 | 9 | 0 | 100 | 0 | 105 |  | 100 | 105 | 0 |  |  | no |
| ***C. ulcerans* s.s.** | 105 |  |  | 105 | 100 | 105 | 0 | 100 | 0 | 9 |  | 100 | 9 | 0 |  |  | yes |

**Table 5.** Database cross-check using the main spectra projections (MSPs) of the *Corynebacterium diphtheriae s*pecies complex (*Cd*SC) from the commercial MBT_K_ as samples, identified by using only the *Cd*SC-MSPs from the MALDI-UP catalogue in order to scrutinise the recent taxonomy (cf. Table 1); bv: biovar; gv: genomovar.

| Reference entry name (MSP) from MBTK | Result with CdSC-MSP from MALDI-UP database<br>**1. hit (score);**<br>**2. hit (score)** |
| --- | --- |
| *Corynebacterium diphtheriae* DSM 44123T<br>DSM_2 | *C. diphtheriae* gravis KL0240 CVUAS (2.68);<br>*C. diphtheriae* gravis KL0167 CVUAS (2.67) |
| *Corynebacterium diphtheriae* CCUG 15935<br>CCUG | *C. diphtheriae* gravis KL0163 CVUAS (2.52);<br>*C. diphtheriae* mitis KL0235 CVUAS (2.50) |
| *Corynebacterium diphtheriae* CCUG 55537<br>CCUG | *C. diphtheriae* mitis KL0252 CVUAS (2.60);<br>*C. diphtheriae* diphtheriae DSM 44123 CVUAS (2.59) |
| *Corynebacterium diphtheriae* DSM 43988 DSM | *C. diphtheriae* diphtheriae DSM 44123 CVUAS (2.67);<br>*C. diphtheriae* gravis KL0163 CVUAS (2.62) |
| *Corynebacterium diphtheriae* 1G12072319_3e<br>MVD | *C. diphtheriae* gravis KL0167 CVUAS (2.63);<br>*C. diphtheriae* gravis KL0237 CVUAS (2.61) |
| *Corynebacterium diphtheriae* 44_R9 ISB | ***C. belfantii* DSM 105776 CVUAS (2.41);**<br>***C. belfantii* KL0171 CVUAS (2.36)** |
| *Corynebacterium diphtheriae* 50_R5 ISB | *C. diphtheriae* gravis KL0163 CVUAS (2.07);<br>*C. diphtheriae* mitis KL0235 CVUAS (2.07) |
| *Corynebacterium diphtheriae* R1_36_29 ISB | *C. diphtheriae* diphtheriae DSM 44123 CVUAS (2.56);<br>*C. diphtheriae* mitis KL0252 CVUAS (2.52) |
| *Corynebacterium pseudotuberculosis* DSM<br>20689T DSM | *C. pseudotuberculosis* bv Ovis DSM 20689 CVUAS (2.48);<br>*C. pseudotuberculosis* bv Equi NTTB 992 CVUAS (2.230) |
| *Corynebacterium pseudotuberculosis* 59_D6_coll<br>ISB | *C. pseudotuberculosis* bv Ovis gv-camelid CVUAS 5583.2 CVUAS<br>(2.53);<br>*C. pseudotuberculosis* bv Ovis gv-camelid CVUAS 32689 CVUAS<br>(2.47) |
| *Corynebacterium pseudotuberculosis* 62_D6_coll<br>ISB | *C. pseudotuberculosis* bv Ovis gv Camelid CVUAS 5583.2 CVUAS<br>(2.36);<br>*C. pseudotuberculosis* bv Ovis gv Camelid CVUAS 32689 CVUAS<br>(2.31) |
| *Corynebacterium pseudotuberculosis* 64_D6_coll<br>ISB | *C. pseudotuberculosis* bv Ovis gv Camelid CVUAS 5583.2 CVUAS<br>(2.37);<br>*C. pseudotuberculosis* bv Ovis DSM 20689 CVUAS (2.29) |
| *Corynebacterium pseudotuberculosis* GD7 GDD | *C. pseudotuberculosis* bv Ovis gv Camelid CVUAS 5583.2 CVUAS<br>(2.54);<br>*C. pseudotuberculosis* bv Ovis gv Camelid CVUAS 32689 CVUAS<br>(2.47) |
| *Corynebacterium pseudotuberculosis* GD8 GDD | *C. pseudotuberculosis* bv Ovis DSM 20689 CVUAS (2.51);<br>*C. pseudotuberculosis* bv Ovis gv Camelid CVUAS 5583.2 CVUAS<br>(2.48) |
| *Corynebacterium pseudotuberculosis*<br>x_x_102968_C IBS | *C. pseudotuberculosis* bv Ovis DSM 20689 CVUAS (2.24);<br>*C. pseudotuberculosis* bv Ovis gv Camelid VUAS 5583.2 CVUAS<br>(2.22) |
| *Corynebacterium ulcerans* DSM 46325T DSM | ***C. ramonii* CVUAS 34267 CVUAS (2.45);**<br>***C. ulcerans* NT 131011719 CVUAS (2.32)** |
| *Corynebacterium ulcerans* 66_D6_coll ISB | *C. ulcerans* NT 141001548 CVUAS (2.12);<br>*C. ulcerans* 07UVF148 LLBB (2.12) |
| *Corynebacterium ulcerans* 67_D6_coll ISB | ***C. ramonii* CVUAS 34267 CVUAS (2.27);**<br>***C. ulcerans* NT 131011719 CVUAS (2.21)** |
| *Corynebacterium ulcerans* 71_D6_coll ISB | *C. ulcerans* DSM 46325 CVUAS (2.25);<br>*C. ulcerans* NT 141001548 CVUAS (2.21) |
| *Corynebacterium ulcerans* MCW_10006 MCW | ***C. ramonii* CVUAS 34267 CVUAS (2.64);**<br>***C. ramonii* CIP 112226 CVUAS (2.61)** |
| *Corynebacterium ulcerans* VA22435_07 ERL | ***C. ramonii* CVUAS 34267 CVUAS (2.41);**<br>***C. ramonii* CIP 112226 CVUAS (2.38)** |

This cross-check investigation revealed that representatives of the new *Cd*SC species may be masked among the earlier reference entries in the commercial MBT_K_ Bruker database.

Consequently, a combination of the in-house extension and a selectively reduced commercial databases creating a modified MBT_K_ was utilised, whereby all MSPs of *C. ulcerans* (s.l.) and *C. diphtheriae* (s.l.) were excluded from the commercial MBT_K_, in order to eliminate any potential inaccuracies in classification (Table 5).

The single spectra identified as *C. ulcerans* s.l. using the MBT_K_, were browsed for the single signal at *m/z* 5,405.4 described by Crestani et al. (2023) as a unique signal marker for *C. ramonii*. In addition to the type strain, seven isolates exhibited a positive signal for this marker (see Fig. 2a, Fig. 2b). The aforementioned conspicuous isolates were subsequently analysed in detail by whole genome sequencing (WGS), following the approach described by Hiller et al. (2024). This analysis assigned the wildlife isolates C86, C104, C139, C167, CVUAS 34267 originating from hedgehogs, and the human isolates KL3299A and CCM 2823 to *C. ramonii*. Once the above mentioned isolates had been reassigned, the two databases were subsequently validated.

### Results with the custom MBT_K_*+MUP database

The custom set of reference spectra from the MUP database (Figure 1), and the application of the described specific *m/z* values, refined and improved the identification results for each species. Thus, using the MBT_K_*+MUP database, all spectra from *C. diphtheriae* s.s. isolates were correctly assigned to the corresponding species (31 out of 31). In comparison to use of the MBT_K_ database, the mean score for this set of spectra could be increased from 2.19 to 2.45. Isolates reliably assigned to *C. belfantii* (type strain and several isolates previously identified as *C. diphtheriae* bv. Belfanti at the LGL Bayern) were correctly identified at species level with the new database, using the first two hit approach. Furthermore, all *C. rouxii* spectra (n=8) were also correctly identified with the MBT_K_*+MUP database (Table 4). Nevertheless, the available number of individual spectra for a formal validation of *C. belfantii* and *C. rouxii* was not sufficient to fulfil the strict formal criteria of the guideline used (BVL, 2022).

Similarly, all spectra of *C. ulcerans* s.s. isolated from a broad collection of host species were correctly assigned with the modified custom MBT_K_*+MUP database. The 88 isolates of *C. silvaticum*, corresponding to the former “wild-boar cluster” of *C. ulcerans* (Rau et al., 2019), were also identified clearly and without discrepancies to *C. ulcerans* or *C. ramonii* with the updated and modified database. In contrast to *C. ulcerans* s.s. and *C. silvaticum*, the limited number of isolates available for *C. ramonii* allows only an initial assessment and no clear differentiation from *C. ulcerans* s.s. (Table S1) using the Biotyper algorithm alone (Compass software, Bruker). The two species, *C. ulcerans* s.s. and *C. ramonii*, were therefore combined in the formal validation step (Table S4f; Table 4). Only the subsequent evaluation of the species-specific *m/z* value of 5,404.5 for *C. ramonii* described by Crestani et al. (2023) allows a separation from *C. ulcerans* s.s.. All isolates’ spectra showing the respective signal were confirmed as closely related to *C. ramonii* by WGS analysis. Remarkably, also strain CCM 2823 (= CIP 54.69) was assigned to *C. ramonii*. A formal validation in the sense of the guideline is not yet conclusive for this step due to the small number of isolates of *C. ramonii* available so far (n=8). This shortage will be remedied as soon as a sufficient number of isolates is available (Crestani et al., 2023; Shitada et al, 2024; Lowe et al., 2025).

Regarding *C. pseudotuberculosis*, the use of the modified and extended MBT_K_*-MUP database shows no divergence of the number of correct assignments compared to the identification using the commercial MBT_K_ database alone. However, a moderate higher mean score value was obtained for the validation spectra set used (2.36 vs. 2.46). In the same way, no false-positive identifications were observed for other species of the genus *Corynebacterium* (n=37), other Actinomycetota species (n=57), or even more distantly related microorganisms (n=750) included in the validation dataset (Table 2). This also applies to the commercial MBT_K_ database.

In conclusion, *C. diphtheriae, C. pseudotuberculosis, C. silvaticum,* and *C. ulcerans* can be reported as validated parameters in our accredited laboratories for official veterinary diagnostics. This achievement is based on the fulfilment of all requirements of the extended reference spectra collection (MBT_K_ + MUP) (BVL, 2022) and the validation procedure specified formal criteria of the BVL technical guidelines. For *C. belfantii, C. ramonii* and *C. rouxii*, the prospects of differentiation are promising, although the number of available spectra for isolates of these species is still too small for a complete formal validation.

For all the seven species of the *Cd*SC, we propose identification hints using the commercial MBT_K_ database to guide the user and to avoid erroneous interpretation of the results (Table 3). In case of the CVUAS, now the custom MBT_K_*+MUP database is used in the regular bacteriological workflow, based on the results of the formal validation (e.g. Walter et al., 2025).

Using the MBT_K_*+MUP database, the species identification of species not belonging to the *Cd*SC is enhanced significantly (Table 2 and 4). In this case, the identification rate of the control group increases from 47.3% for all isolate spectra (n=1,529) using the commercial MBT_K_ to 76.6 % using the custom MBT_K_*+MUP database (e.g. Table S3e vs. Table S4e). Therefore, we conclude that regular database updates are necessary to ensure a comprehensive, efficient and up-to-date database.

As previously demonstrated, the user-driven exchange of custom entries accessible on MUP serves as an effective approach for extending the possibilities of in-house identifications, complementing the periodic updates to the commercial database (Rau et al., 2025). The availability of open access catalogues, like the MALDI User Platform MUP, facilitates the inter-laboratory exchange of spectra. Moreover, the option of downloading spectra collections, such as the MUP or the RKI database, enables users to access and provide data for a broad scientific community (Lasch et al., 2025; CVUAS, 2015). Finally, successful validation can be performed based on a broad collection of well-documented MALDI-TOF MS single spectra to reduce the workload for each single laboratory (BVL, 2022; Rau et al., 2025).

## Conclusions and Prospects

MALDI-TOF MS has become an indispensable tool for a fast and reliable identification of bacteria at the species level in many diagnostic laboratories. However, a sophisticated and up-to-date database is necessary for the exact identification of closely related bacterial species according to current taxonomy, as here demonstrated for the valid species of the *Cd*SC. Identification of bacteria according to the current taxonomy is the basis of treatment of human and animal infections, risk assessment and the implementation of preventive measures such as vaccination programs and ultimately meets the One Health approach.

Here we show, how to keep up with the current taxonomic assignments of bacteria by modifying and extending a current commercial database with own and exchanged MSPs via the globally accessible MALDI-UP catalog for user created reference and validation spectra. The validation process used fulfills the required number of isolates in accordance with the applied guidelines (BVL 2022). This approach of identification of also closely related bacterial species using MALDI-TOF MS is certainly also a model for other bacterial species of interest to react on current taxonomic revisions in a short time frame.

## CRediT authorship contribution statement

Jörg Rau: Supervision, Writing – review & editing, Writing – original draft, Investigation, Formal analysis, Data curation, Methodology, Conceptualisation; Anja Berger: Writing – review & editing, Resources; Alexandra Dangel: Resources, review & editing; Martin Dyk: Investigation, Formal analysis, Data curation, Resources, Methodology; Tobias Eisenberg: Writing – review & editing, Resources; Ekkehard Hiller: Writing – review & editing, Investigation, Formal analysis, Data curation, Methodology; Christiane Hoffmann: Writing – review & editing, Resources; Peter Kutzer: Writing – review & editing, Resources; An Martel: Writing – review & editing, Resources; Andreas Sing: Writing – review & editing; Reinhard Sting: Supervision, Writing – review & editing, Writing – original draft, Resources.

## Funding

Supported by the Bavarian State Ministry of Health, Care, and Prevention and by the German Federal Ministry of Health through the Robert Koch Institute and its National Reference Laboratories Network (grant no. FKZ 1369-359) to the German National Consiliary Laboratory for Diphtheria.

## Acknowledgements

We thank Vanessa Nowak, Sabine Lohrer, Jasmin Scholz, Marion Lindermayer, Anne Könitzer, Wolfgang Schmidt, Helga Kocak, Juliane Breitenberger, Andrea Seifarth, Juliana Webersberger and Turgut-Cengiz Dedeoglu for expert technical assistance.

## Supplemental Material

***Table S1***

Isolates and spectra used.

Data extract from MALDI User Platform (MUP; version 10.02.2025), with metadata for all isolates used in this study and with technical metadata for the individual MALDI-TOF mass spectra used.

---

***Table S2***

List of reference genomes for ANI downloaded from NCBI.

***Table S3a-S2i***

Reports of validations for *Corynebacterium (C.)* identification of species from the *C. diphtheriae* species complex with MALDI-TOF MS using the MALDI Biotyper MBT_K_.

***Table S3a***

Validation for *Corynebacterium* (*C.*) *belfantii* identification with MALDI-TOF MS using the MALDI Biotyper MBT_K_ – Report.

Quantitative evaluation of the MALDI Biotyper, database version K (MBT_K_; Bruker Daltonics) classification performance: results for all single spectra from *C. belfantii* strains and for the non-parameter (i.e. non-*C. belfantii*) isolate set, following the principles of the Guidelines for Validating Species Identifications for Targeted Parameters (BVL, 2022).

***Table S3b***

Validation for *Corynebacterium* (*C.*) *diphtheriae* s.s. identification using the MALDI Biotyper MBT_K_ – Report.

Quantitative evaluation of the MALDI Biotyper, database version K (MBT_K_; Bruker Daltonics) classification performance: results for all single spectra from *C. diphtheriae* s.s. strains and for the non-parameter (i.e. non-*C. diphtheriae* s.s.) isolate set, following the principles of the Guidelines for Validating Species Identifications for Targeted Parameters (BVL, 2022).

***Table S3c***

Validation for *Corynebacterium* (*C.*) *rouxii* identification using the MALDI Biotyper MBT_K_ – Report.

Quantitative evaluation of the MALDI Biotyper, database version K (MBT_K_; Bruker Daltonics) classification performance: results for all single spectra from *C. rouxii* strains and for the non-parameter (i.e. non-*C. rouxii*.) isolate set, following the principles of the Guidelines for Validating Species Identifications for targeted parameters (BVL, 2022).

***Table S3d***

Validation for *Corynebacterium* (*C.*) *diphtheriae* s.l. identification using the MALDI Biotyper MBT_K_ – Report.

Quantitative evaluation of the MALDI Biotyper, database version K (MBT_K_; Bruker Daltonics) classification performance: results for all single spectra from *C. diphtheriae* s.l. strains and for the non-parameter (i.e. non-*C. diphtheriae* s.l.) isolate set, following the principles of the Guidelines for Validating Species Identifications for targeted parameters (BVL, 2022).

***Table S3e***

Validation for *Corynebacterium* (*C.*) *pseudotuberculosis* identification using the MALDI Biotyper MBT_K_ – Report.

Quantitative evaluation of the MALDI Biotyper, database version K (MBT_K_; Bruker Daltonics) classification performance: results for all single spectra from *C. pseudotuberculosis* strains and for the non-parameter (i.e. non-*C. pseudotuberculosis*) isolate set, following the principles of the Guidelines for Validating Species Identifications for targeted parameters (BVL, 2022).

***Table S3f***

Validation for *Corynebacterium* (*C.*) *silvaticum* identification with MALDI-TOF MS using the MALDI Biotyper MBT_K_ – Report.

Quantitative evaluation of the MALDI Biotyper, database version K (MBT_K_; Bruker Daltonics) classification performance: results for all single spectra from *C. silvaticum* strains and for the non-parameter (i.e. non-*C. silvaticum*) isolate set, following the principles of the Guidelines for Validating Species Identifications for targeted parameters (BVL, 2022).

***Table S3g***

Validation for *Corynebacterium* (*C.*) *ramonii* identification with MALDI-TOF MS using the MALDI Biotyper MBT_K_ - Report.

Quantitative evaluation of the MALDI Biotyper, database version K (MBT_K_; Bruker Daltonics) classification performance: results for all single spectra from *C. ramonii* strains and for the non-parameter (i.e. non-*C. ramonii*) isolate set, following the principles of the Guidelines for Validating Species Identifications for targeted parameters (BVL, 2022).

***Table S3h***

Validation for *Corynebacterium* (*C.*) *ulcerans* identification with MALDI-TOF MS using the MALDI Biotyper MBT_K_ - Report.

Quantitative evaluation of the MALDI Biotyper, database version K (MBT_K_; Bruker Daltonics) classification performance: results for all single spectra from *C. ulcerans* strains and for the non-parameter (i.e. non-*C. ulcerans*) isolate set, following the principles of the Guidelines for Validating Species Identifications for targeted parameters (BVL, 2022).

***Table S3i***

Validation for *Corynebacterium* (*C.*) *ulcerans* s.l. identification using the MALDI Biotyper MBT_K_ – Report.

Quantitative evaluation of the MALDI Biotyper, database version K (MBT_K_; Bruker Daltonics) classification performance: results for all single spectra from *C. ulcerans* s.l. strains and for the non-parameter (i.e. non-*C. ulcerans* s.l.) isolate set, following the principles of the Guidelines for Validating Species Identifications for targeted parameters (BVL, 2022).

***Table S4a-S4f***

Reports of validations for *Corynebacterium (C.)* identification of species from the *C. diphtheriae **s***pecies complex with MALDI-TOF MS using the modified MALDI Biotyper MBT_K_* combined with the MALDI-UP extension.

***Table S4a***

Validation of Co*rynebacterium* (*C.*) *belfantii* identification using the modified MALDI Biotyper MBT_K_* combined with the MALDI-UP extension – Report.

Quantitative evaluation of the MALDI Biotyper combined with a database extension (MBT_K_*+ MUP) classification performance: results for all single spectra from *C. belfantii* strains and for the non-parameter (i.e. non-*C. belfantii*) isolate set, following the principles of the Guidelines for Validating Species Identifications for targeted parameters (BVL, 2022).

***Table S4b***

Validation of Co*rynebacterium* (*C.*) *diphtheriae* s.s. identification using the modified MALDI Biotyper MBT_K_* combined with the MALDI-UP extension - Report.

Quantitative evaluation of the MALDI Biotyper combined with a database extension (MBT_K_*+ MUP) classification performance: results for all single spectra from *C. diphtheriae* s.s. strains and for the non-parameter (i.e. non-*C. diphtheriae s.s.*) isolate set, following the principles of the Guidelines for Validating Species Identifications for targeted parameters (BVL, 2022).

***Table S4c***

Validation of Co*rynebacterium* (*C.*) *rouxii* identification using the modified MALDI Biotyper MBT_K_ *combined with the MALDI-UP extension – Report.

Quantitative evaluation of the MALDI Biotyper combined with a database extension (MBT_K_*+ MUP) classification performance: results for all single spectra from *C. rouxii* strains and for the non-parameter (i.e. non-*C. rouxii*) isolate set, following the principles of the Guidelines for Validating Species Identifications for targeted parameters (BVL, 2022).

***Table S4d***

Validation of Co*rynebacterium* (*C.*) *pseudotuberculosis* identification using the modified MALDI Biotyper MBT_K_* combined with the MALDI-UP extension - Report.

Quantitative evaluation of the MALDI Biotyper combined with a database extension (MBT_K_*+ MUP) classification performance: results for all single spectra from *C. pseudotuberculosis* strains and for the non-parameter (i.e. non-*C. pseudotuberculosis*) isolate set, following the principles of the Guidelines for Validating Species Identifications for targeted parameters (BVL, 2022).

***Table S4e***

Validation of Co*rynebacterium* (*C.*) *silvaticum* identification using the modified MALDI Biotyper MBT_K_* combined with the MALDI-UP extension - Report.

Quantitative evaluation of the MALDI Biotyper combined with a database extension (MBT_K_*+ MUP) classification performance: results for all single spectra from *C. silvaticum* strains and for the non-parameter (i.e. non-*C. silvaticum*) isolate set, following the principles of the Guidelines for Validating Species Identifications for targeted parameters (BVL, 2022).

***Table S4f***

Validation of Co*rynebacterium* (*C.*) *ulcerans-ramonii* identification using the modified MALDI Biotyper MBT_K_* combined with the MALDI-UP extension - Report.

Quantitative evaluation of the MALDI Biotyper combined with a database extension (MBT_K_*+ MUP) classification performance: results for all single spectra from *C. ulcerans-ramonii* strains and for the non-parameter (i.e. non-*C. ulcerans-ramonii*) isolate set, following the principles of the Guidelines for Validating Species Identifications for targeted parameters (BVL, 2022).

**Table S5**

Examples of pathogen – host combinations for the species of the *Corynebacterium* (*C.*) *diphtheriae* species complex. References including cases and reviews from 1995 to today.

**Figure S6**

Heatmap showing the average nucleotide identity (ANI) analysis of 184 *Corynebacterium* isolates

## Notes

### Competing Interest Statement

The authors have declared no competing interest.

## References

Aquino de Sá Mda C, Gouveia GV, Krewer Cda C, Veschi JL, de Mattos-Guaraldi AL, da Costa MM. 2013. Distribution of PLD and FagA, B, C and D genes in *Corynebacterium pseudotuberculosis* isolates from sheep and goats with caseus lymphadenitis. Genet Mol Biol. 36:265–268. 10.1590/S1415-47572013005000013.

Badell E, Hennart M, Rodrigues C, Passet V, Dazas M, Panunzi L, Bouchez V, Carmi–Leroy A, Toubiana J, Brisse S. 2020. *Corynebacterium rouxii* sp. nov., a novel member of the diphtheriae species complex. Research in Microbiology. 171:122–127. 10.1016/j.resmic.2020.02.003.

Badenschier F, Berger A, Dangel A, Sprenger A, Hobmaier B, Sievers C, Prins H, Dörre A, Wagner - Wiening C, Küper-Schiek W, Wichmann O, Sing A. 2022).Outbreak of imported diphtheria with *Corynebacterium diphtheriae* among migrants arriving in Germany, 2022. Eurosurveillance. 27:46. https://www.eurosurveillance.org/content/10.2807/1560-7917.ES.2022.27.46.2200849.

Barksdale L Linder R, Sulea IT, Pollice M. 1981. Phospholipase D activity of *Corynebacterium pseudotuberculosis* (*Corynebacterium ovis*) and *Corynebacterium ulcerans*, a distinctive marker within the genus *Corynebacterium*. J Clin Microbiol. 13:335–343. 10.1128/jcm.13.2.335-343.1981.

Bartlett A, Padfield D, Lear L, Bendall R, Vos M. 2022. A comprehensive list of bacterial pathogens infecting humans. Microbiology. 168:12. 10.1099/mic.0.001269.

Berger A, Dangel A, Peters M, Mühldorfer K, Braune S, Eisenberg T, Szentiks CA, Rau J, Konrad R, Hörmansdorfer S, Ackermann N, Sing A. 2019. Tox-positive *Corynebacterium ulcerans* in hedgehogs, Germany. Emerging Microbes and Infection. 8:211–217. 10.1080/22221751.2018.1562312.

Berger A, Dangel A, Melnikov VG, Bengs K, Rupp T, Mappes HJ, Schneider C, Sing A. 2025. Human infections by novel zoonotic species *Corynebacterium silvaticum*, Germany. Emerging Infectious Disease 31:1450–1454. 10.3201/eid3107.250086.

Brengenzer T, Frei R, Ohnacker H, Zimmerli W. 1997. *Corynebacterium pseudotuberculosis* infection in a butcher. Clinical Microbiology and Infection. 3:696–698. 10.1111/j.1469-0691.1997.tb00482.x.

Bruker. 2018. Product information - Release Notes - MBT Compass Library Revision E (2018).

Bruker. 2022. Product information - Release Notes - MBT Compass Library Revision K (2022).

BVL. 2022. Federal Office of Consumer Protection and Food Safety: Guidelines for validating species identifications using matrix-assisted laser desorption/ionisation time-of-flight mass spectrometry (MALDI-TOF-MS) in a single laboratory or in laboratory networks, pp. 1–32 https://www.bvl.bund.de/SharedDocs/Downloads/07_Untersuchungen/Guidelines_for_validating_species_identifications_using_MALDI-TOF-MS.pdf;jsessionid=385A5C363D8718FF3D44B0A04694F886.2_cid372?blob=publicationFile&v=4.

CVUAS. 2015. Chemisches und Veterinäruntersuchungsamt Stuttgart: MALDI-TOF MS user platform MALDI-UP. Retrieved from https://www.maldi-up.ua-bw.de, Accessed 10th Feb 2025.

Crestani C, Arcari G, Landier A, Passet V, Garnier D, Brémont S, Armatys N, Carmi-Leroy A, Toubiana J, Badell E, Brisse S. 2023. *Corynebacterium ramonii* sp. nov., a novel toxigenic member of the *Corynebacterium diphtheriae* species complex. Res Microbiol. 174:104113. 10.1016/j.resmic.2023.104113.

Contzen M, Sting R, Blazey B, Rau J. 2011. *Corynebacterium ulcerans* from diseased wild boars carrying *Corynebacterium diphtheriae*-like tox genes. Zoon Publ Health. 58:479–488c. https://onlinelibrary.wiley.com/doi/abs/10.1111/j.1863-2378.2011.01396.x.

Dangel A, Berger A, Rau J, Eisenberg T, Kämpfer P, Margos G, Contzen M, Busse H-J, Konrad R, Peters M, Sting R, Sing A. 2020. *Corynebacterium silvaticum* sp. nov., a unique group of NTTB corynebacteria in wild boar and roe deer. Int J Syst Evol Microbiol. 70:3614–3624. 10.1099/ijsem.0.004195.

Dazas M, Badell E, Carmi-Leroy A, Criscuolo A, Brisse S. 2018. Taxonomic status of *Corynebacterium diphtheriae biovar* Belfanti and proposal of *Corynebacterium belfantii* sp. nov. Int J Syst Evol Microbiol. 68:3826–3831. 10.1099/ijsem.0.003069.

Deneke C, Brendebach,H, Uelze L, Borowiak M, Malorny B, Tausch SH. 2021. Species-Specific Quality Control, Assembly and Contamination Detection in Microbial Isolate Sequences with AQUAMIS. Genes. 12:644. 10.3390/genes12050644.

Domenis L, Spedicato R, Orusa R, Goria M, Sant S, Guidetti C, Robetto S. 2017. The diagnostic activity on wild animals through the description of a model case report (caseous lymphadenitis by *Corynebacterium pseudotuberculosis* associated with *Pasteurella* spp and parasites infection in an alpine ibex –Capra ibex). Open Veterinary Journal. 7:384–390. 10.4314/ovj.v7i4.15.

Domenis L, Spedicato R, Pepe E, Orusa R, Robetto S. 2018. Caseous Lymphadenitis caused by *Corynebacterium pseudotuberculosis* in Alpine Chamois (*Rupicapra r. rupicapra*): a review of 98 cases. Journal of Comparative Pathology. 161:11–19. 10.1016/j.jcpa.2018.04.003.

Eberson FA.1918. Bacteriologic study of the diphtheroid organisms with special reference to Hodgkin’s disease. Journal of Infectious Diseases 23:1–42. 10.1086/infdis/23.1.1.

Eisenberg T, Contzen M, Kutzer P, Peters M, Sing A, Rau J. 2014. Non toxigenic tox-bearing *Corynebacterium ulcerans* infection in game in Germany. Emerging Infection Diseases. 20:448–452. 10.3201/eid2003.130423.

Eisenberg T, Mauder N, Contzen M, Rau J, Ewers C, Schlez K, Althoff G, Schauerte N, Geiger C, Margos G, Konrad R, Sing A. 2015. Outbreak with clonally related isolates of *Corynebacterium ulcerans* in a group of water rats. BMC Microbiology. 15:42. 10.1186/s12866-015-0384-x.

Engler KH, Glushkevich T, Mazurova IK, George RC, Efstratiou A. 1997. A modified Elek test for detection of toxigenic corynebacteria in the diagnostic laboratory. J Clin Microbiol, 35:495–498. 10.1128/jcm.35.2.495-498.1997.

European Centre for Disease Prevention and Control (ECDC). 2022. Rapid Risk Assessment: Increase of reported diphtheria cases due to *Corynebacterium diphtheriae* among migrants in Europe – 6 October 2022. Stockholm. https://www.ecdc.europa.eu/sites/default/files/documents/diphtheria-cases-migrants-europe-corynebacterium-diphtheriae-2022.pdf.

European Centre for Disease Prevention and Control (ECDC). 2024. Diphtheria. In: ECDC. Annual epidemiological report for 2022. Stockholm. https://www.ecdc.europa.eu/sites/default/files/documents/DIPH_AER_2022_Report.pdf.

EU-Commission. 2018. COMMISSION IMPLEMENTING DECISION (EU) 2018/945 of 22 June 2018 on the communicable diseases and related special health issues to be covered by epidemiological surveillance as well as relevant case definitions. Official Journal of the European Union. L170/1 6.7.2018. https://eur-lex.europa.eu/eli/dec_impl/2018/945/oj/eng.

Gant MS, Chamot-Rooke J. 2024. Present and future perspectives on mass spectrometry for clinical microbiology. Microbes Infect. 26:105296. 10.1016/j.micinf.2024.105296.

Grönthal TSC, Lehto AK, Aarnio SS, Eskola EK, Aimo-Koivisto EM, Karlsson T, Koskinen HI, Barkoff AM, He Q, Lienemann T, Rimhanen-Finne R, Mykkänen A. 2024. Pastern dermatitis outbreak associated with toxigenic and non-toxigenic Corynebacterium diphtheriae and non-toxigenic Corynebacterium ulcerans at a horse stable in Finland, 2021. Zoon Pub Health. 71:127-135. 10.1111/zph.13090.

Goris J, Konstantinidis KT, Klappenbach JA, Coenye T, Vandamme P, Tiedje JM. 2007. DNA-DNA hybridization values and their relationship to whole-genome sequence similarities. Int J Syst Evol Microbiol. 57(Pt 1):81–91. Epub 2007/01/16. 10.1099/ijs.0.64483-0.

Hacker E, Antunes CA, Mattos-Guaraldi AL, Burkovski A, Tauch A. 2016. *Corynebacterium ulcerans*, an emerging human pathogen. Future Microbiology. 11:1191–1208. 10.2217/fmb-2016-0085.

Hennart M, Crestani C, Bridel S, Armatys N, Brémont S, Carmi-Leroy A, Landier A, Passet V, Fonteneau L, Vaux S, Toubiana J, Badell E, Brisse S. 2023. A global C*orynebacterium diphtheriae* genomic framework sheds light on current diphtheria reemergence. Peer Community Journal. 3:e76. 10.24072/pcjournal.307.

Hillan A, Gibbs T, Weaire-Buchanan G, Brown T, Peng S, McEvoy SP, Parker E. 2023. Zoonotic transmission of diphtheria toxin-producing *Corynebacterium ulcerans*. Zoon Pub Health. 71:157–169. 10.1111/zph.13094.

Hiller E, Hörz V, Sting R. 2024. *Corynebacterium pseudotuberculosis*: Whole genome sequencing reveals unforeseen and relevant genetic diversity in this pathogen. PLOS one. 0309282. 10.1371/journal.pone.0309282.

Hoefer A, Seth-Smith Helena, Palma F, Schindler S, Freschi L, Dangel A, Berger A, D’Aeth J, Indra A, Fry NK, Palm D, Sing A, Brisse S, Egli, A. 2025. *Corynebacterium diphtheriae* outbreak in migrant populations in Europe. N Engl J Med 392:2334–234510.1056/NEJMoa2311981.

Jacquinet S, Martini H, Mangion JP, Neusy S, Detollenaere A, Hammami N, Bruggeman L, Hoorelbeke B, Pierard D, Cornelissen L. 2023. Outbreak of *Corynebacterium diphtheriae* among asylum seekers in Belgium in 2022: operational challenges and lessons learnt. Euro Surveill. 28:2300130. 10.2807/1560-7917.ES.2023.28.44.2300130.

Kostrzewa M, Maier T. 2017. Criteria for development of MALDI-TOF mass spectral database in H.N. Shah, S.E. Gharbia (Eds.), MALDI-TOF and Tandem MS for clinical microbiology, John Wiley & Sons Ltd., Hoboken, USA 10.1002/9781118960226.ch2.

Lasch P, Nattermann H, Erhard H, Stämmler M, Grunow R, Bannert N, Appel B, Naumann D. 2008. MALDI-TOF mass spectrometry compatible inactivation method for highly pathogenic microbial cells and spores. Analytical Chemistry. 80:2026–2034. 10.1021/ac701822j.

Lasch P, Beyer W, Bosch A, Borriss R, Drevinec M, Dupke S, Ehling-Schulz M, Gao X, Grunow R, Jacob D, Klee SR, Paauw A, Rau J, Schneider A, Scholz HC, Stämmler M, Tam LTT, Tomaso H, Werner G, Döllinger J. 2025. A MALDI-ToF mass spectrometry database for identification and classification of highly pathogenic bacteria. Nature Scientific Data. 12:187. 10.1038/s41597-025-04504-z.

Lehmann KB, Neumann R. 1886. Atlas und Grundriss der Bakteriologie und Lehrbuch der speziellen bakteriologischen Diagnostik. 1st ed., Munich, Germany.

Lowe CF, Ritchie G, Crestani C, Imperial M, Matic N, Payne M, Stefanovic A, Diana Whellams D, Brisse S, G. Romney MG. 2025. Detection of diphtheria toxin-producing *Corynebacterium ramonii* in wounds of an urban inner-city population in Vancouver, Canada, 2019-2023. Emerg Infect Dis. 31:323–327. 10.3201/eid3102.241472.

Malorny B, Scheel K, Rau J, Beyer W, Buschulte A, Nöckler K, Kreienbrock L. 2020. Onlineumfrage zur Anwendung von molekularbiologischen Typisierungsverfahren und MALDI-TOF-MS in diagnostischen Laboren in Deutschland. Journal of Consumer Protection and Food Safety. 15:387–391. 10.1007/s00003-020-01297-8.

Martel A, Boyen F, Rau J, Eisenberg T, Sing A, Berger A, Chiers K, Van Praet S, Verbanck S, Vervaeke M, Pasmans F. 2021. Widespread disease in hedgehogs (*Erinaceus europaeus*) caused by toxigenic *Corynebacterium ulcerans*. Emerg Infect Dis. 27:2686–2690. 10.3201/eid2710.203335.

Meinel DM, Margos G, Konrad R, Krebs S, Blum H, Sing A. 2014. Next generation sequencing analysis of nine *Corynebacterium ulcerans* isolates reveals zoonotic transmission and a novel putative diphtheria toxin encoding pathogenicity island. Genome Med. 6:13. 10.1186/s13073-014-0113-3.

Miers KC, Ley WB. 1980. *Corynebacterium pseudotuberculosis* infection in the horse: study of 117 clinical cases and consideration of etiopathogenesis. J Am Vet Med Assoc. 177:250–253. 10.2460/javma.1980.177.03.250.

Nouioui I, Carro L, García-López M, Meier-Kolthoff JP, Woyke T, Kyrpides NC, Pukall R, Klenk HP, Goodfellow M, Göker M. 2018. Genome-based taxonomic classification of the phylum Actinobacteria. Front Microbiol. 22:9, 2007. 10.3389/fmicb.2018.02007.

Oberreuter H, Dyk M, Rau J. 2023. Validated differentiation of *Listeria monocytogenes* serogroups by FTIR spectroscopy using an Artificial Neural Network based classifier in an accredited official food control laboratory. Clinical Spectroscopy 5:100030. 10.1016/j.clispe.2023.100030.

Oliveira A, Oliveira LC, Aburjaile F, Benevides L, Tiwari S, Jamal SB, Silva A, Figueiredo HCP, Ghosh P, Portela RW, De Carvalho Azevedo VA, Wattam AR. 2017. Insight of genus *Corynebacterium*: Ascertaining the role of pathogenic and non-pathogenic species. Front Microbiol. 8:1937. 10.3389/fmicb.2017.01937.

Otsuji K, Fukuda K, Endo T, Shimizu S, Harayama N, Ogawa M, Yamamoto A, Umeda K, Umata T, Seki H, Iwaki M, Kamochi M, Saito M. 2017. The first fatal case of *Corynebacterium ulcerans* infection in Japan. JMM Case Rep. 4(8):e005106. 10.1099/jmmcr.0.005106.

Peel MM, Palmer GG, Stacpoole AM, Kerr TG. 1997. Human lymphadenitis due to *Corynebacterium pseudotuberculosis:* report of ten cases from Australia and review. Clin Infect Dis. 24:185–91. 10.1093/clinids/24.2.185.

Pranada, BP, Schwarz G, Kostrzewa M. 2016. MALDI Biotyping for microorganism identification in clinical microbiology in R. Cramer (Ed.), Advances in MALDI and Laser-induced soft ionization mass spectrometry, Springer International Publishing, Switzerland pp. 197–222. 10.1007/978-3-319-04819-2_11.

Prates FD, Araújo MRB, Sousa EG, Ramos JN, Viana MVC, Soares SdC, dos Santos LS, Azevedo VAdC. 2024. First pangenome of *Corynebacterium rouxii*, a potentially toxigenic species of *Corynebacterium diphtheriae* Complex. Bacteria. 3:99–117. 10.3390/bacteria3020007.

Prygiel M, Polak M, Mosiej E, Wdowiak K, Formińska K, Zasada AA. 2022. New *Corynebacterium s*pecies with the potential to produce diphtheria toxin. Pathogens. 11:1264. 10.3390/pathogens11111264.

Rau J, Blazey B, Contzen M, Sting R. 2012. *Corynebacterium ulcerans*-Infektion bei einem Reh (*Capreolus capreolus*). Berliner und Münchener Tierärztliche Wochenschrift. 125:159–162. https://vetline.de/abszess-europaeisches-reh-capreolus-capreolus-corynebacterium-ulcerans-ft-ir-tox-gen/150/3130/68865.

Rau J, Eisenberg T, Männig A, Wind C, Lasch P, Sting R. 2016. MALDI-UP – an internet platform for the exchange of mass spectra – user guide for https://maldi-up.ua-bw.de/. Aspects of Food Control and Animal Health. 1:1–17. http://ejournal.cvuas.de/issue201601.asp.

Rau J, Eisenberg T, Peters M, Berger A, Kutzer P, Lassnig H, Hotzel H, Sing A, Sting R, Contzen M. 2019. Reliable differentiation of a non-toxigenic tox gene bearing *Corynebacterium ulcerans* variant frequently isolated from game animals using MALDI-TOF MS. Vet Microbiol. 237:108399. 10.1016/j.vetmic.2019.108399.

Rau J, Eisenberg T, Wind C, Huber I, Pavlovic M, Becker R. 2022. Aus der § 64 LFGB-Arbeitsgruppe MALDI-TOF: Leitlinien für die Validierung von Spezies-Identifizierungen mittels MALDI-TOF-MS. Journal of Consumer Protection and Food Safety, 17:97–101. 10.1007/s00003-021-01353-x.

Rau J, Etter J, Frentzel H, Lasch P, Contzen M. 2025. Reliable delineation of *Bacillus cytotoxicus* from other members of the *Bacillus cereus* group by MALDI-TOF MS – An extensive validation study. Food Control, 167:110825. 10.1016/j.foodcont.2024.110825.

Riegel, P., Ruimy, R., de Briel, D., Prévost, G., Jehl, F, Christen, R., Monteil, H. 1995. Taxonomy of *Corynebacterium diphtheriae* and related taxa, with recognition of *Corynebacterium ulcerans* sp. nov. nom. Rev. FEMS Microbiology Letters. 126:271–276. 10.1016/0378-1097(95)00022-W.

Schlez K, Eisenberg T, Rau J, Dubielzig, S, Kornmayer M, Wolf G, Berger A, Dangel A, Hoffmann C, Ewers C, Sing A. 2021. *Corynebacterium rouxii*, a recently described member of the *C. diphtheriae* group isolated from three dogs with ulcerative skin lesions. Antonie van Leuvenhoek. 114:1361–1371. 10.1007/s10482-021-01605-8.

Selim SA. 2001. Oedematous skin disease of buffaloes in Egypt. Journal of Veterinary Medicine, Series B 48:241–258. 10.1046/j.1439-0450.2001.00451.x.

Shitada C, Moriguchi M, Hayashi H, Matsumoto K, Mori M, Tokuoka E, Yahiro S, Gejima S, Horiba K, Yamamoto T, Takahashi M, Kuroda M. 2024. Genomic analysis of novel bacterial species *Corynebacterium ramonii* ST344 clone strains isolated from human skin ulcer and rescued cats in Japan. Zoonotic Diseases. 4:234–244. 10.3390/zoonoticdis4040020.

Sing A, Konrad R, Meinel DM, Mauder N, Schwabe I, Sting R. 2025. Corynebacterium rouxii in a free-roaming red fox: case report and historical review on diphtheria in animals. Infection. 3/2025. https://www.springermedizin.de/toxigenic-corynebacterium-ulcerans-in-raw-milk-of-a-cow-with-acu/50665762.

Slinko V, Guglielmino C, Uren A, Smith J, Neucom D, Smoll N, Graham R, Fang N, Smith H, Armstrong A, Kenny A, Farmer J, Quagliotto C, Jennison A. 2023. Several confirmed and probable zoonotic cases of toxigenic *Corynebacterium ulcerans*, Queensland, Australia. Communicable Diseases Intelligence 2023:47. 10.33321/cdi.2023.47.53.

Sting R, Rietschel W, Polley B, Süß-Dombrowski C, Rau J. 2017. *Corynebacterium pseudotuberculosis* infection in a dromedary (*Camelus dromedarius*) in Germany. Berliner und Münchener Tierärztliche Wochenschrift. 130:511–516. https://vetline.de/corynebacterium-pseudotuberculosis-infection-in-a-dromedary-camelus-dromedarius-in-germany/150/3130/105368.

Sting R, Geiger C, Rietschel W, Blazey B, Schwabe I, Rau J, Schneider-Bühl L. 2022. *Corynebacterium pseudotuberculosis* Infections in Alpacas (*Vicugna pacos*). Animals. 12:1612. 10.3390/ani12131612.

Sting R, Pölzelbauer C, Eisenberg T, Bonke R, Blazey B, Peters M, Riße K, Sing A, Berger A, Dangel A, Rau J. 2023. *Corynebacterium ulcerans* Infections in Eurasian Beavers (*Castor fiber*). Pathogens. 12:979. 10.3390/pathogens12080979.

Tagini F, Pillonel T, Croxatto A, Bertelli C, Koutsokera A, Lovis A, Greub G. 2018. Distinct genomic features characterize two clades of *Corynebacterium diphtheriae*: Proposal of *Corynebacterium diphtheriae* subsp. *diphtheriae* subsp. nov. and *Corynebacterium diphtheriae* subsp. *lausannense* subsp. Nov.. Front Microbiol. 9:1743. 10.3389/fmicb.2018.01743.

Thomas A, Slifka AM, Hendrickson SM, Amanna IJ, Slifka MK. 2022. Active circulation of *Corynebacterium ulcerans* among nonhuman primates. Microbiol Spectr. 10:e00894–22. 10.1128/spectrum.00894-22.

Thompson JE. 2022. Matrix-assisted laser desorption ionization-time-of-flight mass spectrometry in veterinary medicine: Recent advances (2019-present). Vet World. 15:2623–2657. 10.14202/vetworld.2022.2623-2657.

Timofte D, Overesch G, Spergser J. 2023. MALDI-TOF MS analysis for identification of veterinary pathogens from companion animals and livestock species. In HN Shan, SE Gharbia et al., (Eds). Microbiological identification using MALDI-TOF and tandem mass spectrometry: Industrial and environmental applications. Chapter 12. Wiley. 10.1002/9781119814085.ch12.

Wang H, Wang X, Cao Y, Chen Y, Zou Z, Lu X, Shan F, Tu J, Liu J, Liu J, Sa J, Zhou N, Peng S, Zou J, Shen X, Zhai J, Chen Z, Holmes EC, Chen W, Shen Y0. 2024. Identification of *Corynebacterium ulcerans* and *Erysipelothrix* sp. in Malayan pangolins – a potential threat to public health?. MSphere. 0:e00551-24. 10.1128/MSPhere.00551-24.

Walter B, Lück K, Adam M, Winter B, Rau J, Knab N, Welker A, Aichinger E, Brockmann S. 2025. A cross-sectional study on diphtheria and other wound infections among asylum seekers arriving in Baden-Wuerttemberg, Germany, August – October 2024. Poster. ESCMID 11.-15. April 2025.

## Reference

BVL. 2022. Federal Office of Consumer Protection and Food Safety:. Guidelines for validating species identifications using matrix-assisted laser desorption/ionisation time- of-flight mass spectrometry (MALDI-TOF-MS) in a single laboratory or in laboratory networks, pp. 1–32 https://www.bvl.bund.de/SharedDocs/Downloads/07_Untersuchungen/Guidelines_for_validating_species_identifications_using_MALDI-TOF-MS.pdf;jsessionid=385A5C363D8718FF3D44B0A04694F886.2_cid372?blob=publicationFile&v=4.

