## Supplement 5 Table for "Up-to-date MALDI-TOF MS based identification of the complete *Corynebacterium diphtheriae* species complex for improved diagnostics"

| **Corynebacterium diphtheria s.s.** | | | | | | |
| --- | --- | --- | --- | --- | --- | --- |
| C. diphtheriae s.s. (Lehmann & Neumann, 1896) is the causative agent of classical diphtheria of the upper respiratory tract in humans, a potentially life-threatening, highly transmissible disease. As a consequence of vaccination, the incidence of diphtheria in Europe is currently of minor importance, occurring predominantly in the form of cutaneous lesions. These may be associated with travel to, or origin from regions where the disease is endemic (Badenschier et al., 2022; ECDC, 2022; ECDC, 2024; Hennart et al., 2023; Hoefer et al, 2025; Jaquinet et al., 2023; Walter et al., 2025). The pathological syndrome is associated with the impact of diphtheria toxin (DT), an exoprotein that can be produced by certain types of C. diphtheriae, coded by the corresponding tox gene (Prygiel et al., 2022). Infections in animals are described rarely (Sing et al., 2025a). | | | | | | |
| **Major pathogenicity** | | | |  |  |  |
| ***tox*-gen** | | | **Phospholipase D** |  |  |  |
| **positive** | | **negative** |  |  |  |  |
| **ELEK-test** | |  | **CAMP-test** |  |  |  |
| **positive** | **negative** |  |  |  |  |  |
| **TTB** | **NTTB** | **NT** |  | **disease** | **host** | **References** |
| x | x | x | negative | diphtheria (TTB) | human | ECDC, 2024 |
| **x** |  | **x** |  |  | dog | Sing et al., 2016; Sing et al., 2025 |
| **x** | **x** |  |  |  | cat | Sing et al., 2016; Tyler et al., 2022 |
|  |  | x |  | metritis | pig | Sing et al., 2025a |
| x |  | x |  | Pastern Dermatitis / conjunctivitis draining wound infection  non-healing pyogenic stake wound thoracic wound pus | horse | Museux et al., 2023; Sing et al., 2016; Sing et al., 2025a |

| **Corynebacterium belfantii** | | | | | | |
| --- | --- | --- | --- | --- | --- | --- |
| *C. belfantii* (Dazas et al, 2018) is equivalent to the former biovar Belfanti of *C. diphtheriae*, which was elevated to species rank. This species now also includes the simultaneously described subspecies *C. diphtheriae subsp. lausannense* (Tagini et al., 2018), which is an established human pathogen (Bartlett et al., 2022; Prygiel et al. 2022). | | | | | | |
| **Major pathogenicity** | | | |  |  |  |
| ***tox*-gen** | | | **phospholipase D** |  |  |  |
| **positive** | | **negative** |  |  |  |  |
| **ELEK-test** | |  | **CAMP-test** |  |  |  |
| **positive** | **negative** |  |  |  |  |  |
| **TTB** | **NTTB** | **NT** |  | **disease** | **host** | **References** |
| no | no | **x** | negative | respiratory | human | Dazas et al., 2018 |
|  |  | **x** |  |  | cow | Sing et al., 2025b |

| **Corynebacterium rouxii** | | | | | | |
| --- | --- | --- | --- | --- | --- | --- |
| *C. rouxii* (Badell et al., 2020) was formerly assigned to *C. belfantii.* This species has so far been isolated in the context of skin infections in dogs, cats and humans (Schlez et al., 2021; Bartlett et al., 2022). Some strains bear the *tox*-gene (Prates et al., 2024). | | | | | | |
| **Major pathogenicity** | | | |  |  |  |
| ***tox*-gen** | | | **Phospholipase D** |  |  |  |
| **positive** | | **negative** |  |  |  |  |
| **ELEK-test** | |  | **CAMP-test** |  |  |  |
| **positive** | **negative** |  |  |  |  |  |
| **TTB** | **NTTB** | **NT** |  | **disease** | **host** | **References** |
| no |  | **x** | negative | cutaneous or peritoneum infections | human | Badell et al., 2020 |
|  |  | x | negative | Skin lesion; purulent dermatitis, otitis externa, sore | dog | Badell et al., 2020; Schlez et al., 2022; Museux et al., 2023; Sing et al., 2025 |
|  |  | **x** | negative | Inflammation of mammary gland | red fox | Schlez et al., 2022; Sing et al., 2025 |
|  | x | x | negative | Ulcerative injury, chronic otitis | cat | Museux et al., 2023; Prates et al., 2024 |
|  |  | x |  |  | hedgehog | Martel et. al., 2021; this study |

| **Corynebacterium ulcerans (lineage 1)** | | | | | | |
| --- | --- | --- | --- | --- | --- | --- |
| *C. ulcerans* s.s. (Riegel, 1995) has already been identified as a pathogen in humans and in different mammalian taxa (e.g. Artiodactyla, Perissodactyla, Carnivora, Eulipotyphla, Pholidota, Rodentia, Primates), including exotic, game, pet and livestock animals (Berger et al., 2019; Hillan et al., 2023; Eisenberg et al., 2015; Grönthal et al., 2024; Sting et al., 2023; Martel et al., 2021; Thomas et al., 2022; Wang et al., 2024). In addition to wound infections, *C. ulcerans* also causes diphtheria-like diseases of the upper respiratory tract in humans and has surpassed the incidence of *C. diphtheriae* infections in industrialised nations since several years (ECDC, 2022). In contrast to *C. diphtheriae*, *C. ulcerans* is mainly acquired by humans through contact with animals, especially cats and dogs, and thus represents an important zoonotic pathogen agent (Hillan et al., 2023; Meinel et al., 2014; Otsuji et al., 2017; Slinko et al., 2023). | | | | | | |
| **Major pathogenicity** | | | |  |  |  |
| ***tox*-gen** | | | **Phospholipase D** |  |  |  |
| **positive** | | **negative** |  |  |  |  |
| **ELEK-test** | |  | **CAMP-test** |  |  |  |
| **positive** | **negative** |  |  |  |  |  |
| **TTB** | **NTTB** | **NT** |  | **disease** | **host** | **References** |
| **x** | **x** | **x** | positive | abscess  pneumonia  diphtheria like disease (TTB)  air way obstruction | **human** | Crestani et al., 2023, Museux et al., 2023, Hillan et al., 2023, Wake et al., 2021, Tiwari et al., 2008 |
| **x** |  |  |  |  | Macaques, monkeys | Tiwari et al., 2008 |
| **x** |  |  |  |  | otters | Tiwari et al., 2008 |
|  |  | **x** |  |  | **ferrets** | Marini et al., 2014 |
|  |  | **x** | positive |  | lion | Seto et al., 2008 |
| **x** | **x** | **x** |  | chronic otitis; sores | **dogs, cats** | Crestani et al., 2023; Museux et al., 2023; Shitada et al., 2024 |
|  |  | x | positive |  | **water rats** | Eisenberg et al., 2015 |
| **x** | ? |  |  |  | **rat** | Museux et al., 2023 |
|  | **X** |  |  |  | **rabbit** | Museux et al., 2023 |
| **x** |  |  | positive |  | **beaver** | Sting et al., 2023 |
| x |  |  |  |  | goat | Tiwari et al., 2008 |
| **X** |  |  |  | mastitis | **cows** | Tiwari et al., 2008; Sing et al. 2025 |
| x |  |  |  |  | dromedary | Tiwari et al., 2008 |
| x |  |  | positive |  | killer whale | Seto et al., 2008 |
| **x** |  |  | positive |  | **pig** | Berger et al., 2013; Schuhegger et al., 2009 |
| x |  | **x** |  |  | **donkey, horses** | Crestani et al., 2023; Tiwari et al., 2008 |
|  |  | x |  |  | **pangulins** | Wang et al., 2024 (Clade 1) |
| **x** |  |  |  |  | **Japanese shrew-moles** | Katsukawa et al., 2016 |
| **x** | **x** | **x** | positive | wound infections | **hedgehogs** | Berger et al., 2019; Martel et al., 2021 |
| **x** |  |  |  |  | **ural owl** | Katsukawa et al., 2016 |

| **Corynebacterium ramonii (former C. ulcerans lineage 2)** | | | | | | |
| --- | --- | --- | --- | --- | --- | --- |
| *C. ramonii* (Crestani et al., 2023) was formerly separated from *C. ulcerans*, representing the former genetical lineage 2 and has recently been defined as a novel zoonotic species affecting humans, dogs and cats. Notably, *C. ramonii* has the potential to carry the *tox*-gene (Crestani et al., 2023; Shitada et al., 2024). | | | | | | |
| **Major pathogenicity** | | | |  |  |  |
| ***tox*-gen** | | | **phospholipase D** |  |  |  |
| **positive** | | **negative** |  |  |  |  |
| **ELEK-test** | |  | **CAMP-test** |  |  |  |
| **positive** | **negative** |  |  |  |  |  |
| **TTB** | **NTTB** | **NT** |  | **disease** | **host** | **References** |
| **x** | **x** | **x** | positive | diphtheria like disease (TTB) | **human** | Crestani et al., 2023; Lowe et al., 2025 |
| **x** |  |  |  |  | **dog** | Crestani et al., 2023 |
|  |  | **x** | positive |  | **red fox** | Sting et al., 2015; Schlez et al., 2022 |
|  | **X (Vero cell positive)** |  |  |  | **cat** | Shitada et al., 2024 |
|  |  | **x** |  |  | **hedgehog** | Martel et al., 2021; this study |

| **Corynebacterium silvaticum (former C. ulcerans NTTB wild-boar cluster)** | | | | | | |
| --- | --- | --- | --- | --- | --- | --- |
| *C. silvaticum* (Dangel et al., 2020) was previously classified as a wild boar cluster of *C. ulcerans*. This novel species has been identified as the causative agent of caseous abscesses in wild boars, Iberian pigs, and on the basis of one report in a roe deer (Contzen et al., 2011; Rau et al., 2012; Eisenberg et al., 2014; Viana et al., 2023). Recently, a few cases of human infections via contact with wild boar have been reported (Berger et al., 2025). | | | | | | |
| **Major pathogenicity** | | | |  |  |  |
| ***tox*-gen** | | | **phospholipase D** |  |  |  |
| **positive** | | **negative** |  |  |  |  |
| **ELEK-test** | |  | **CAMP-test** |  |  |  |
| **positive** | **negative** |  |  |  |  |  |
| **TTB** | **NTTB** | **NT** |  | **disease** | **host** | **References** |
| **x** | **x** |  | positive | lymphadenitis | **human** | Berger et al. 2025 |
|  | **x** |  | positive | lymphnode abscess | **wild boar** | Rau et al., 2019; Dangel et al., 2020 |
|  |  |  |  |  | **free ranging pigs** | Viana et al., 2023 |
|  | x |  | positive | lymphnode abscess | **roe deer** | Rau et al., 2012; Dangel et al., 2020 |

| **Corynebacterium pseudotuberculosis** | | | | | | |
| --- | --- | --- | --- | --- | --- | --- |
| *C. pseudotuberculosis* (Eberson, 1918) is the causative agent of chronic infections characterised by formation of abscesses in lymph nodes mainly in sheep, goats, and camelids and is therefore referred to as caseous lymphadenitis (CLA) (Domenis, 2017). The pathogen also causes ulcerative lymphangitis, the so-called pigeon fever in horses. *C. pseudotuberculosis* isolates of water buffaloes are the only isolates known to produce DT and are associated with oedematous skin disease (OSD) (Sting et al., 2017; Sting et al., 2022; Hiller et al., 2024; Selim 2001). Infections with these isolates are rarely encountered in other animals. In general, pseudotuberculosis is a rare zoonosis, which is transmitted via contact with the pathogen, particularly through affected animals (Bregenzer et al., 1997; Peel et al., 1997). Typing of *C. pseudotuberculosis* is traditionally based on testing on nitrate reductase activity, which differentiates isolates into the two biovar Equi (exhibits nitrate reductase activity) and Ovis (lack of nitrate reductase activity). With few exceptions, *C. pseudotuberculosis* isolates originating from horses and water buffaloes belong to the biovar Equi and isolates originating from sheep and goats belong to the biovar Ovis. Furthermore, camelid isolates carrying the nitrate reductase genes, but lacking nitrate reductase activity due to gene mutations could recently be detected. Those isolates were defined as the novel type biovar Ovis, genomovar Camelid, and show a close genetic relationship to biovar Equi isolates (Hiller et al., 2024). | | | | | | |
| **Major pathogenicity** | | | |  |  |  |
| ***tox*-gen** | | | **phospholipase D** |  |  |  |
| **positive** | | **negative** |  |  |  |  |
| **ELEK-test** | |  | **CAMP-test** |  |  |  |
| **positive** | **negative** |  |  |  |  |  |
| **TTB** | **NTTB** | **NT** |  | **disease** | **host** | **References** |
| **biovar Ovis** | | | | | | |
| - | - | x | positive | necrotizing lymphadenitis/ pneumonia | (human) | Heggelund et al., 2015; Peel et al., 1997 |
|  |  | x | positive | caseous lyphadenitis (CLA) | **Sheep, goat** | Hiller et al., 2024 |
|  |  | x | positive |  | (w**ildebeest)**  (**llama**)  **alpine ibex**  **alpine chamois**  **huemul**  **roe deer**  **antelope** | Viana et al., 2017; Hiller et al., 2024; Müller et al., 2011; Domenis et al., 2018; Domenis et al., 2017;  Di Donato et al., 2024 |
|  |  |  | positive | ulcerative granulomatous lesions and mastitis | **cattle** | Sing et al., 2025b |
|  |  | x | positive |  | (camel)  (alpine chamois) | Viana et al., 2017  Domenis et al., 2018 |
| **biovar Equi** | | | | | | |
|  |  |  |  |  | **sheep** | Hiller et al., 2024 |
|  |  |  | positive | ulcerative lymphangitis, pigeon fever | **horse** | Hiller et al., 2024 |
| **x** |  | x | positive | Oedematous skin disease (OSD) | **water-buffalo**  **cattle** | Hiller et al., 2024; Viana et al., 2017; Selim 2001 |
| **biovar Ovis / genomovar Camelid** | | | | | | |
|  |  | x | positive | caseous lyphadenitis (CLA) | **camelids** | Sting et al., 2017; Sting et al., 2022; Hiller et al., 2024 |

References

Badenschier F, Berger A, Dangel A, Sprenger A, Hobmaier B, Sievers C, Prins H, Dörre A, Wagner -Wiening C, Küper-Schiek W, Wichmann O, Sing A. 2022).Outbreak of imported diphtheria with *Corynebacterium diphtheriae* among migrants arriving in Germany, 2022. Eurosurveillance. 27:46. <https://www.eurosurveillance.org/content/10.2807/1560-7917.ES.2022.27.46.2200849>.

Badell E, Hennart M, Rodrigues C, Passet V, Dazas M, Panunzi L, Bouchez V, Carmi–Leroy A, Toubiana J, Brisse S. 2020. Corynebacterium rouxii sp. nov., a novel member of the diphtheriae species complex. Research in Microbiology. 171:122-127. <https://doi.org/10.1016/j.resmic.2020.02.003>.

Bartlett A, Padfield D, Lear L, Bendall R, Vos M. 2022. A comprehensive list of bacterial pathogens infecting humans. Microbiology. 168:12. <https://doi.org/10.1099/mic.0.001269>.

Berger A, [Bo](https://onlinelibrary.wiley.com/authored-by/Boschert/V.)s[chert](https://onlinelibrary.wiley.com/authored-by/Boschert/V.) V, [K](https://onlinelibrary.wiley.com/authored-by/Konrad/R.)o[nrad](https://onlinelibrary.wiley.com/authored-by/Konrad/R.) R, Schmidt-Wieland T, [Hörmansdorfer](https://onlinelibrary.wiley.com/authored-by/Hörmansdorfer/S.) S, Eddicks M, Sing A. 2013. Two Cases of cutaneous diphtheria associated with occupational pig contact in Germany. Zoonoses and Public Health. 60:539-542. <https://doi.org/10.1111/zph.12031>.

Berger A, Dangel A, Peters M, Mühldorfer K, Braune S, Eisenberg T, Szentiks CA, Rau J, Konrad R, Hörmansdorfer S, Ackermann N, Sing A. 2019. Tox-positive Corynebacterium ulcerans in hedgehogs, Germany. Emerging Microbes and Infection. 8:211-217. <https://doi.org/10.1080/22221751.2018.1562312>.

Berger A, Dangel A, Melnikov VG, Bengs K, Rupp T, Mappes HJ, Schneider C, Sing A. 2025. Human infections by novel zoonotic species Corynebacterium silvaticum, Germany. Emerging Infectious Disease 31:1450-1454.

<https://doi.org/10.3201/eid3107.250086>.

Brengenzer T, Frei R, Ohnacker H, Zimmerli W. 1997. *Corynebacterium pseudotuberculosis* infection in a butcher. Clinical Microbiology and Infection. 3:696-698. https://doi.org/10.1111/j.1469-0691.1997.tb00482.x.

Crestani C, Arcari G, Landier A, Passet V, Garnier D, Brémont S, Armatys N, Carmi-Leroy A, Toubiana J, Badell E, Brisse S. 2023. Corynebacterium ramonii sp. nov., a novel toxigenic member of the Corynebacterium diphtheriae species complex. Res Microbiol. 174:104113. <https://doi.org/10.1016/j.resmic.2023.104113>.

Contzen M, Sting R, Blazey B, Rau J. 2011. *Corynebacterium ulcerans* from diseased wild boars carrying *Corynebacterium diphtheriae*-like tox genes. Zoon. Publ. Health. 58:479-488c. <https://onlinelibrary.wiley.com/doi/abs/10.1111/j.1863-2378.2011.01396.x>.

Dangel A, Berger A, Rau J, Eisenberg T, Kämpfer P, Margos G, Contzen M, Busse H-J, Konrad R, Peters M, Sting R, Sing A. 2020. Corynebacterium silvaticum sp. nov., a unique group of NTTB corynebacteria in wild boar and roe deer. Int J Syst Evol Microbiol. 70:3614-3624. https://doi.org/10.1099/ijsem.0.004195.

Dazas M, Badell E, Carmi-Leroy A, Criscuolo A, Brisse S. 2018. Taxonomic status of Corynebacterium diphtheriae biovar Belfanti and proposal of Corynebacterium belfantii sp. nov. Int J Syst Evol Microbiol. 68:3826-3831. <https://doi.org/10.1099/ijsem.0.003069>.

Di Donato A, Gambi L, Ravaioli V, Perulli S, Cirasella L, Rossini R, Luppi A, Tosi G, Fiorentini L. (2024). First report of caseous lymphadenitis by Corynebacterium pseudotubercolosis and pulmonary verminosis in a roe deer (Capreolus capreolus Linnaeus, 1758) in Italy. Animals. 14:566. https://doi.org/10.3390/ani14040566.

Heggelund L, Gaustad P, Håvelsrud OE, Blom J, Borgen L, Sundset A, Sørum H, Frøland SS. 2015. *Corynebacterium pseudotuberculosis* pneumonia in a veterinary student infected during laboratory work. Open Forum Infect Dis. 2:ofv053. https://doi.org/ 10.1093/ofid/ofv053.

Hennart M, Crestani C, Bridel S, Armatys N, Brémont S, Carmi-Leroy A, Landier A, Passet V, Fonteneau L, Vaux S, Toubiana J, Badell E, Brisse S. 2023. A global C*orynebacterium diphtheriae* genomic framework sheds light on current diphtheria reemergence. Peer Community Journal. 3:e76. <https://doi.org/10.24072/pcjournal.307>.

Hillan A, Gibbs T, Weaire-Buchanan G, Brown T, Peng S, McEvoy SP, Parker E. 2023. Zoonotic transmission of diphtheria toxin-producing *Corynebacterium ulcerans.* Zoonoses and Public Health. 71:157-169. <https://doi.org/10.1111/zph.13094>.

Hiller E, Hörz V, Sting R. 2024. *Corynebacterium pseudotuberculosis*: Whole genome sequencing reveals unforeseen and relevant genetic diversity in this pathogen. PLOS one. 0309282. <https://doi.org/10.1371/journal.pone.0309282>.

Hoefer A, Seth-Smith Helena, Palma F, Schindler S, Freschi L, Dangel A, Berger A, D’Aeth J, Indra A, Fry NK, Palm D, Sing A, Brisse S, Egli, A. 2025. *Corynebacterium diphtheriae* outbreak in migrant populations in Europe. N Engl J Med 392:2334-2345 <https://doi.org/10.1056/NEJMoa2311981>.

Jacquinet S, Martini H, Mangion JP, Neusy S, Detollenaere A, Hammami N, Bruggeman L, Hoorelbeke B, Pierard D, Cornelissen L. 2023. Outbreak of *Corynebacterium diphtheriae* among asylum seekers in Belgium in 2022: operational challenges and lessons learnt. Euro Surveill. 28:2300130. <https://doi.org/10.2807/1560-7917.ES.2023.28.44.2300130>.

Katsukawa C, Umeda K, Inamori I, Kosono Y, Tanigawa T, Komiya T, Iwaki M, Yamamoto A, Nakatsu S. 2016. Toxigenic *Corynebacterium ulcerans* isolated from a wild bird (ural owl) and its feed (shrew-moles): comparison of molecular types with human isolates. BMC Res Notes. 9:181. https://doi.org/10.1186/s13104-016-1979-5.

Kelly EJ, Rood, KA, Skirpstunas R. 2012. Abscesses in captive Elk associated with *Corynebacterium pseudotuberculosis,* Utah, USA. Journal of Wildlife Diseases 48:803-805. <https://doi.org/10.7589/0090-3558-48.3.803>

Lehmann KB, Neumann R. 1886. Atlas und Grundriss der Bakteriologie und Lehrbuch der speziellen bakteriologischen Diagnostik. 1st ed., Munich, Germany.

Lowe CF, Ritchie G, Crestani C , Imperial M, Matic N, Payne M, Stefanovic A, Diana Whellams D, Brisse S, G. Romney MG. 2025. Detection of diphtheria toxin-producing *Corynebacterium ramonii* in wounds of an urban inner-city population in Vancouver, Canada, 2019-2023. Emerging Infectious Diseases. 31:323-327. https://doi.org/10.3201/eid3102.241472.

Marini RP, Cassiday PK, Venezia J, Shen Z, Buckley EM, Peters Y, et al. 2014. Corynebacterium ulcerans in ferrets. Emerg Infect Dis. 20:159-161. <https://doi.org/10.3201/eid2001.130675>.

Martel A, Boyen F, Rau J, Eisenberg T, Sing A, Berger A, Chiers K, Van Praet S, Verbanck S, Vervaeke M, Pasmans F. 2021. Widespread disease in hedgehogs (Erinaceus europaeus) caused by toxigenic Corynebacterium ulcerans. Emerg Infect Dis. 27:2686-2690. <https://doi.org/10.3201/eid2710.203335>.

Meinel DM, Margos G, Konrad R, Krebs S, Blum H, Sing A. 2014. Next generation sequencing analysis of nine Corynebacterium ulcerans isolates reveals zoonotic transmission and a novel putative diphtheria toxin encoding pathogenicity island. Genome Med. 6:13. <https://doi.org/10.1186/s13073-014-0113-3>.

Museux K, Arcari G, Rodrigo G, Hennart M, Badell E, Toubiana J, Brisse S. 2023. Corynebacteria of the *diphtheriae* species complex in companion animals: Clinical and microbiological cCharacterization of 64 cases from France. Microbiol Spectr. 11:e0000623. <https://doi.org/10.1128/spectrum.00006-23>.

Müller B, de Klerk-Lorist LM, Henton MM, Lane E, Parsons S, van Pittius NCG, Kotze A, an Helden PD, Tanner M. 2011. Mixed infections of *Corynebacterium pseudotuberculosis* and non-tuberculous mycobacteria in South African antelopes presenting with tuberculosis-like lesions. Veterinary microbiology. 147:340-345. https://doi.org/10.1016/j.vetmic.2010.07.017.

Schuhegger R, Schoerner C, Dlugaiczyk J, Lichtenfeld I, Trouillier A, Zeller-Peronnet V, et al. 2009. Pigs as source for toxigenic Corynebacterium ulcerans. Emerg Infect Dis. 15:1314-1315. <https://doi.org/10.3201/eid1508.081568>.

Selim SA. 2001. Oedematous skin disease of buffaloes in Egypt. Journal of Veterinary Medicine, Series B 48:241-258. <https://doi.org/10.1046/j.1439-0450.2001.00451.x>.

Seto Y, Komiya T, Iwaki M, Kohda T, Mukamoto M, Takahashi M, et al. 2008. Properties of corynephage attachment site and molecular epidemiology of Corynebacterium ulcerans isolated from humans and animals in Japan. Jpn J Infect Dis. 61:116–22. <https://doi.org/10.7883/yoken.JJID.2008.116>.

Shitada C, Moriguchi M, Hayashi H, Matsumoto K, Mori M, Tokuoka E, Yahiro S, Gejima S, Horiba K, Yamamoto T, Takahashi M, Kuroda M. 2024. Genomic analysis of novel bacterial species *Corynebacterium ramonii* ST344 clone strains isolated from human skin ulcer and rescued cats in Japan. Zoonotic Diseases. 4:234-244. <https://doi.org/10.3390/zoonoticdis4040020>.

Sing A, Konrad R, Meinel DM, Mauder N, Schwabe I, Sting R. 2016. *Corynebacterium diphtheriae* in a free-roaming red fox: case report and historical review on diphtheria in animals. Infection. 44:441-445. <https://link.springer.com/article/10.1007/s15010-015-0846-y>.

Sing A, Konrad R, Meinel DM, Mauder N, Schwabe I, Sting R. 2025a. *Corynebacterium rouxii* in a free-roaming red fox: case report and historical review on diphtheria in animals. Infection. 3/2025. https://www.springermedizin.de/toxigenic-corynebacterium-ulcerans-in-raw-milk-of-a-cow-with-acu/50665762.

Sing A, Luaces LM, Dangel A, Deramo S, Bengs K, Melnikov VG, Berger A. 2025b. Toxigenic *Corynebacterium ulcerans* in raw milk of a cow with acute mastitis: case report and historical review on milk-transmitted diphtheria. Infection. 53:767–774. <https://doi.org/10.1007/s15010-025-02477-0>.

Slinko V, Guglielmino C, Uren A, Smith J, Neucom D, Smoll N, Graham R, Fang N,Smith H, Armstrong A, Kenny A, Farmer J, Quagliotto C, Jennison A. 2023. Several confirmed and probable zoonotic cases of toxigenic *Corynebacterium ulcerans*, Queensland, Australia. Communicable diseases intelligence 2023:47. <https://doi.org/10.33321/cdi.2023.47.53>.

Sting R, Ketterer-Pintur S, Contzen M, Mauder N, Sing A, Süß-Dombrowski C. 2015. Toxigenic *Corynebacterium ulcerans* isolated from a free-roaming red fox (*Vulpes vulpes*). Berliner und Münchener Tierärztliche Wochenschrift. 128: 04-208. <https://vetline.de/toxigenic-corynebacterium-ulcerans-isolated-from-a-free-roaming-red-fox-vulpes-vulpes/150/3130/87702>.

Sting R, Rietschel W, Polley B, Süß-Dombrowski C, Rau J. 2017. *Corynebacterium pseudotuberculosis* infection in a dromedary (*Camelus dromedarius*) in Germany. Berliner und Münchener Tierärztliche Wochenschrift. 130:511-516.

<https://vetline.de/corynebacterium-pseudotuberculosis-infection-in-a-dromedary-camelus-dromedarius-in-germany/150/3130/105368>.

Sting R, Geiger C, Rietschel W, Blazey B, Schwabe I, Rau J, Schneider-Bühl L. 2022. Corynebacterium pseudotuberculosis infections in alpacas (Vicugna pacos). Animals. 12:1612. <https://doi.org/10.3390/ani12131612>.

Sting R, Pölzelbauer C, Eisenberg T, Bonke R, Blazey B, Peters M, Riße K, Sing A, Berger A, Dangel A, Rau J. 2023. *Corynebacterium ulcerans* infections in Eurasian beavers (*Castor fiber*). Pathogens. 12:979. <https://doi.org/10.3390/pathogens12080979>.

Tagini F, Pillonel T, Croxatto A, Bertelli C, Koutsokera A, Lovis A, Greub G. 2018. Distinct genomic features characterize to clades of Corynebacterium diphtheriae: Proposal of *Corynebacterium diphtheriae* sbsp. *diphtheriae* subsp. nov. and *Corynebacterium diphtheriae* subsp. *lausannense* subsp. nov. Front Microbiol. 9:1743. <https://doi.org/10.3389/fmicb.2018.01743>.

Thomas A, Slifka AM, Hendrickson SM, Amanna IJ, Slifka MK. 2022. Active Circulation of *Corynebacterium ulcerans* among nonhuman primates. Microbiol Spectr10:e00894-22. <https://doi.org/10.1128/spectrum.00894-22>.

Tiwari TSP, Golaz A, Yu DT, Ehresmann KR, Jones TF, Hill HE, Cassiday PK, Pawloski LC, Moran JS, Popovic T, Wharton M,2008. Investigations of 2 cases of diphtheria-lke iIllness due to toxigenic Corynebacterium ulcerans. Clinical Infectious Diseases, 46:395–401. <https://doi.org/10.1086/525262>.

Tyler R Jr, Rincon L, Weigand MR, Xiaoli L, Acosta AM, Kurien D, Ju H, Lingsweiler S, Prot EY. 2022. Toxigenic *Corynebacterium diphtheriae* infection in cat, Texas, USA. Emerg Infect Dis. 28:1686-1688. https://doi.org/10.3201/eid2808.220018.

Viana MVC, Figueiredo H, Ramos R, Guimarães LC, Pereira FL, Dorella FA, Selim SAK, Salaheldean M, Silva A, Wattam AR, Azevedo V. 2017. Comparative genomic analysis between *Corynebacterium pseudotuberculosis* strains isolated from buffalo. PLoS One. 2(4):e0176347. <https://doi.org/10.1371/journal.pone.0176347>.

Viana MVC, Galdino JH, Profeta R, Oliveira M, Tavares L, de Castro Soares S, Carneiro P, Wattam AR, Azevedo V. 2023. Analysis of *Corynebacterium silvaticum* genomes from Portugal reveals a single cluster and a clade suggested to produce diphtheria toxin. PeerJ. 11:e14895. <https://doi.org/10.7717/peerj.14895>.

Wake K, Kikuchi J, Uchida M, Nemoto M, Kaji Y, Yokoyama T, Suzuki H, Fukushima A, Yamamoto A, Iwaki M, Ono K. 2021. Transmission of toxigenic *Corynebacterium ulcerans* infection with airway obstruction from cats to a human. Acute Med Surg 12;8(1):e705. <https://doi.org/10.1002/ams2.705>.

Walter B, Lück K, Adam M, Winter B, Rau J, Knab N, Welker A, Aichinger E, Brockmann S. 2025. A cross-sectional study on diphtheria and other wound infections among asylum seekers arriving in Baden-Wuerttemberg, Germany, August – October 2024. Poster. ESCMID 11.-15. April 2025.
