## Supplementary figures and images for "Up-to-date MALDI-TOF MS based identification of the complete *Corynebacterium diphtheriae* species complex for improved diagnostics"

### Supplement 6 Figure

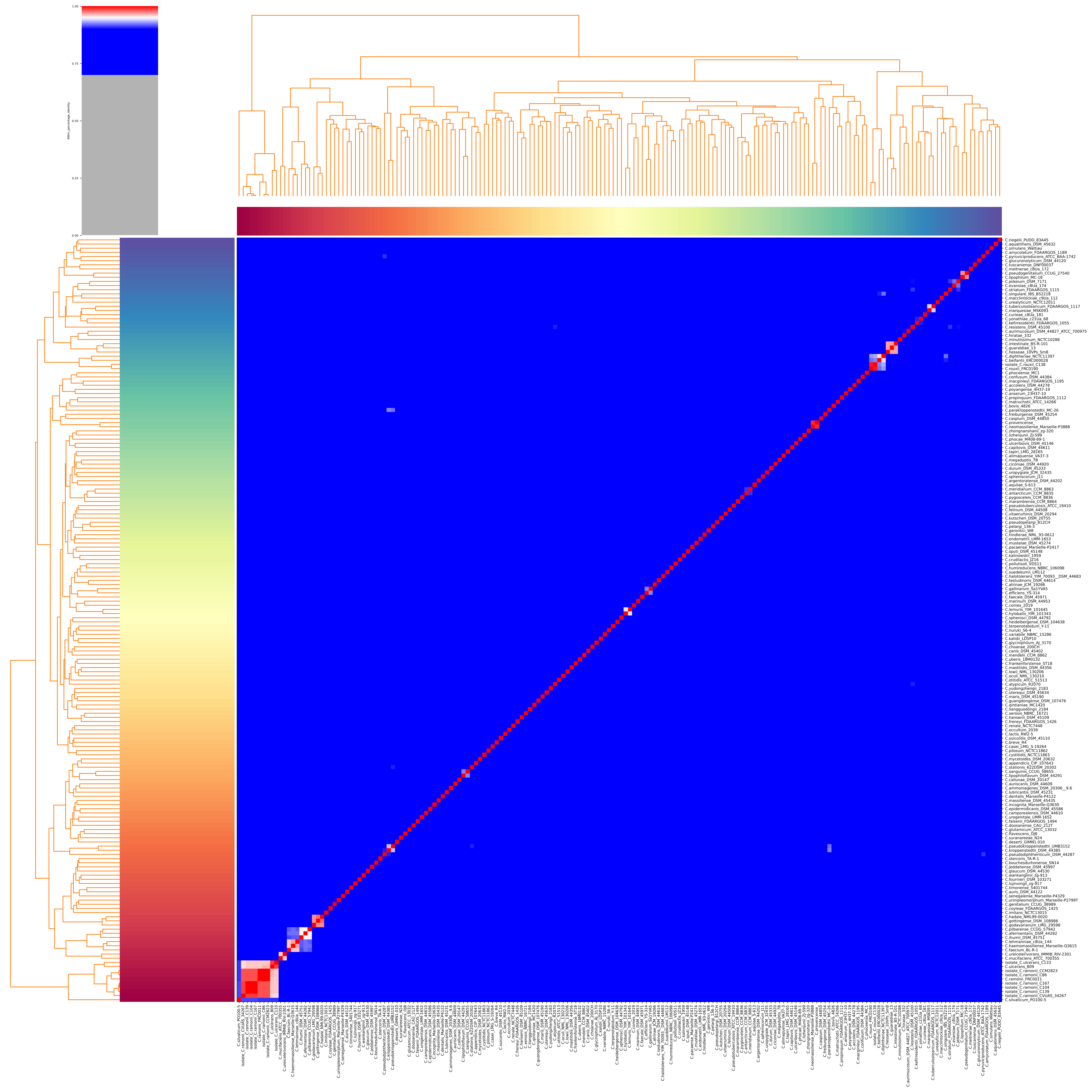
